# High-dimensional geometry of population responses in visual cortex

**DOI:** 10.1101/374090

**Authors:** Carsen Stringer, Marius Pachitariu, Nicholas Steinmetz, Matteo Carandini, Kenneth D. Harris

## Abstract

A neuronal population encodes information most efficiently when its activity is uncorrelated and high-dimensional, and most robustly when its activity is correlated and lower-dimensional. Here, we analyzed the correlation structure of natural image coding, in large visual cortical populations recorded from awake mice. Evoked population activity was high dimensional, with correlations obeying an unexpected power-law: the *n^th^* principal component variance scaled as 1/*n*. This was not inherited from the 1/*f* spectrum of natural images, because it persisted after stimulus whitening. We proved mathematically that the variance spectrum must decay at least this fast if a population code is smooth, i.e. if small changes in input cannot dominate population activity. The theory also predicts larger power-law exponents for lower-dimensional stimulus ensembles, which we validated experimentally. These results suggest that coding smoothness represents a fundamental constraint governing correlations in neural population codes.

## Introduction

The visual cortex contains millions of neurons, and the patterns of activity that images evoke in these neurons form a “population code”. The structure of this code is largely unknown, due to the lack of techniques able to record from large populations. Nonetheless, the population code is the subject of long-standing theories.

Two extreme alternatives of such theories are the “efficient coding hypothesis” and the “columnar hypothesis”. Efficient coding^1–3^ maintains that neural codes maximize information transmission by eliminating correlations present in natural image inputs. Such codes are highdimensional, which can allow complex features to be read out by simple downstream networks^4–6^. In contrast, the columnar hypothesis holds that all neurons in a cortical column encode similar information^7^; the resulting lowdimensional codes have been suggested to allow reliable computations to arise from inherently noisy circuits^8^. In between these two extremes lie a range of possibilities, which can be characterized by their dimensionality.

Several experimental studies have suggested that neural codes are confined to low-dimensional planes^9–17^. Nevertheless, theoretical considerations show that such results are inevitable given stimuli or tasks of limited complexity^18^: the responses to a set of *n* stimuli, for example, have to lie on an *n* — 1 dimensional plane. The dimensionality of the cortical code thus remains an open question, which can only be answered by recording the responses of large numbers of neurons to large numbers of stimuli.

Here, we recorded the simultaneous activity of ~ 10,000 neurons in mouse visual cortex, in response to thousands of natural images. Our results were consistent with neither the efficient coding hypothesis nor the columnar hypothesis: responses were not confined to a low-dimensional plane, but neither was the code uncorrelated. Instead, responses occupied a multidimensional space with the variance in the *n^th^* dimension scaling as a power law *n*^−*α*^, where *α* ≈ 1. This power-law scaling did not reflect correlations in the images themselves, as it persisted when showing decorrelated images. Instead, we hypothesized it arises from smoothness constraints. We show mathematically that if variances decay slower than a power law with exponent *α* = 1 + 2/*d*, where *d* is the dimension of the input ensemble, then the space of neural activity must be fractal, i.e. show increasingly rough structure at finer and finer scales. We verified that variances are almost as large as allowed by this bound by presenting stimulus ensembles of varying dimension. These findings suggest that the population code of visual cortex is determined by two constraints: efficiency, to make best use of limited numbers of neurons, and smoothness, which allows similar images to evoke similar responses.

### Simultaneous recordings of ~10,000 neurons

To obtain simultaneous recordings of ~ 10,000 cells from mouse VI, we employed resonance-scanning two-photon calcium microscopy, using 11 imaging planes spaced at 35*ı*m (Fig. 1a). The slow timecourse of the GCaMP6s sensor allowed activity to be detected at a 2.5 Hz scan rate, and an efficient data processing pipeline^19^ allowed large numbers of cells to be detected accurately (Fig. 1b). Natural image scenes (Imagenet database^20^) were presented on an array of 3 monitors surrounding the mouse (Fig. 1c), at an average of 1 image/s. Cells were tuned to the natural image stimuli: in experiments in which responses to 32 images were averaged over 96 repeats (Fig. 1d), stimulus responses accounted for 55.4±3.3% (SE, n=4 recordings) of the trial-averaged variance. Consistent with prior reports^21–23^, neuronal responses were sparse: only a small fraction (13.4=L·1.0% SE, n=4 recordings) of cells were driven more than two standard deviations above their baseline firing rate by any particular stimulus.

**Fig. 1.**
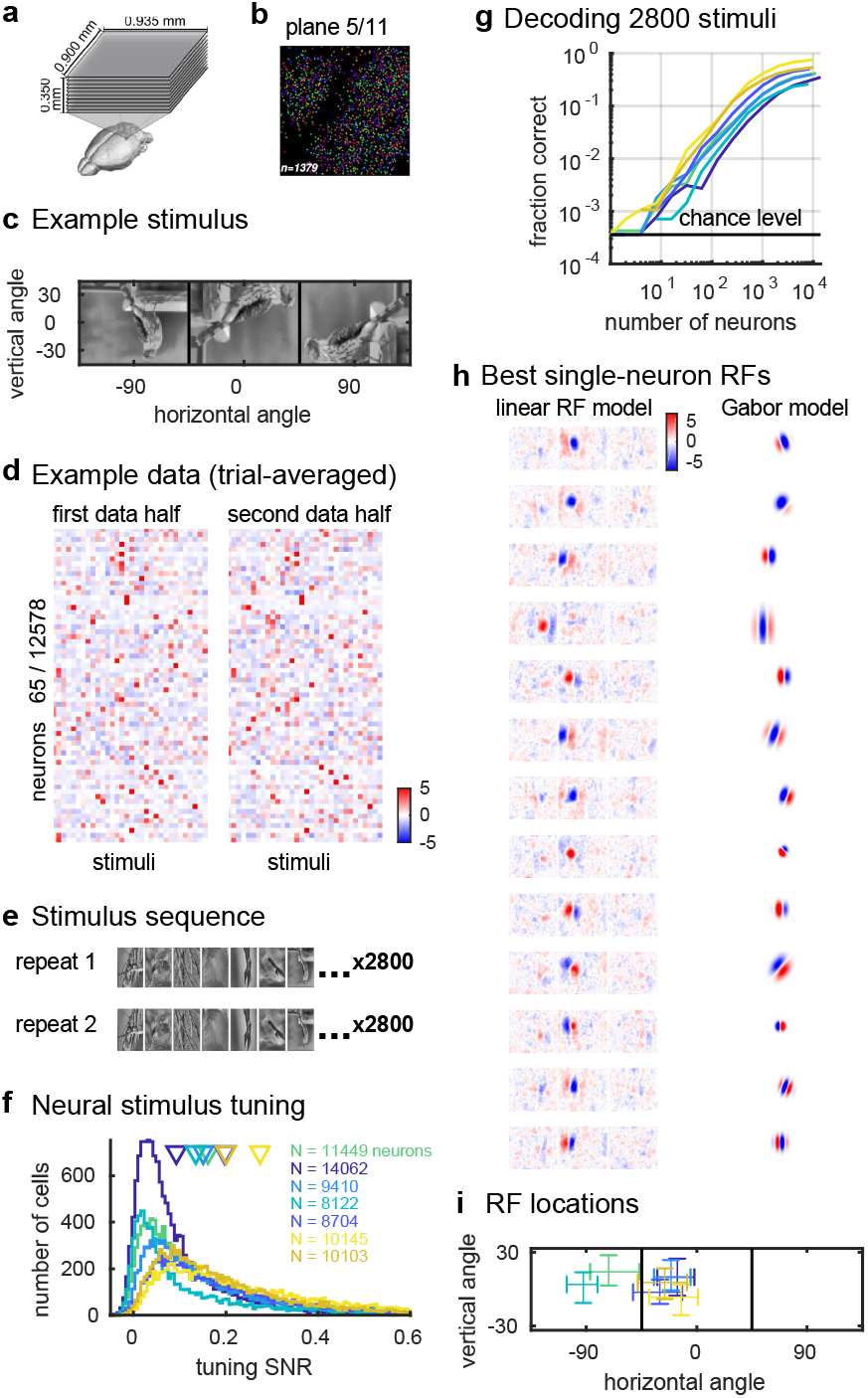
Population coding of visual stimuli. **a**, Simultaneous recording of ~ 10,000 neurons using 11-plane two-photon calcium imaging, **b,** Randomly-pseudocolored cells in an example imaging plane, **c,** Example natural image stimulus spans three screens surrounding the mouse’s head, **d,** Mean responses of 65 randomly-chosen neurons to 32 image stimuli (96 repeats), **e,** To allow cross-validated analysis of responses to a large image ensemble, a sequence of 2800 stimuli was repeated twice during the recording, **f,** Distribution of single-cell signal-to-noise ratios (SNR) (2800 stimuli, two repeats). Colors denote recordings; arrows represent means, **g,** Stimulus decoding accuracy as a function of neuron count for each recording, **h,** Receptive fields (RFs) fit using reduced-rank regression or Gabor models, for best-fit neurons, **i,** Distribution of the receptive field centers, plotted on the left and center screens (line denotes screen boundary). Each cross represents a different recording, with error bars representing 95% confidence intervals on that recording’s mean RF position.

For our main experiments, we presented a sequence of 2,800 image stimuli repeated in succession. Two repeats were used to allow the maximum number of images to be analyzed, while still allowing analyses based on cross-validation (Fig. 1e). A majority of neurons (81.4=L·5.1% SE, n=7 recordings) showed correlation between repeats at *p* < 0.05; Extended Data Fig. 1a,b). Nevertheless, consistent with previous reports^24^, singletrial responses showed substantial trial-to-trial variability. Cross-validation showed that stimulus responses accounted for on average 13.9=L·1.7% of the single-trial variance (Extended Data Fig. 1c), and the average signal-to-noise ratio was 17.3±2.4% (Fig. 1f). This level of trial-to-trial variability was not due our particular recording method: measuring responses to the same stimuli electrophysiologically yielded a similar signal-to-noise ratio (Extended Data Fig. 2). Despite trial-to-trial variability, however, the activity recorded on a single trial from the 10,000 cell populations contained substantial information about the sensory stimuli. Indeed, a simple nearest-neighbor decoder, trained on one repeat and tested on the other, was able to identify the presented stimulus with up to 75.5% accuracy (Fig. 1g; range 25.4%-75.5%; median 41.7% compared to chance level of 0.036%, n=7 recordings). Decoding accuracy did not appear to have saturated at population size 10,000, suggesting that performance would further increase with more neurons.

Neurons had similar visual properties to previous reports^23,25^, and their responses were only partially captured by classical linear-nonlinear models, consistent with a vast literature indicating that natural image responses in VI are are only partially approximated by classical RF models^26–30^. We calculated a receptive field (RF) for each cell from its responses to natural images in two ways: by fitting linear RFs regularized with a reduced rank method; or by searching for an optimal Gabor filter that was rectified/quadrature filtered to simulate classical simple/complex cell responses. As expected from retinotopy, the RF locations of simultaneously recorded neurons overlapped but there was a high diversity of receptive field sizes and shapes (Fig. 1h; Extended Data Fig. 3, Extended Data Fig. 4). Both RF models, however, explained less than 20% of the stimulus-related variance (the linear model explained 11.4±0.7% SE, and the Gabor model explained 18.5=L·1.0% SE, n=7 recordings each).

### Dimensionality and power-law scaling of variances

To characterize the geometry of the visual cortical population code, we developed a method of cross-validated principal component analysis (PC A). PC A provides for each integer *n* the maximum fraction of variance that can be accounted for by linear variations along an optimal *n*-dimensional subspace. Direct application of PCA to neural data, however, would measure the dimensionality of both the stimulus representation and trial-to-trial fluctuations. Since the present study is concerned only with the former, we developed a cross-validated approach that focuses only on stimulus representation (Fig. 2a), and also projected out dimensions corresponding to ongoing activity (see Extended Data Fig. 5; see Methods). To estimate the amount of stimulus-related variance in an optimal *n*-dimensional plane, we chose this plane by PCA of one stimulus repeat (the training set), and measured the fraction of the second repeat’s variance that was confined to this plane. We confirmed that this technique can recover the true variances using simulations of neural data with the same noise statistics as our recordings (Extended Data Fig. 6; Supplementary Information).

**Fig. 2.**
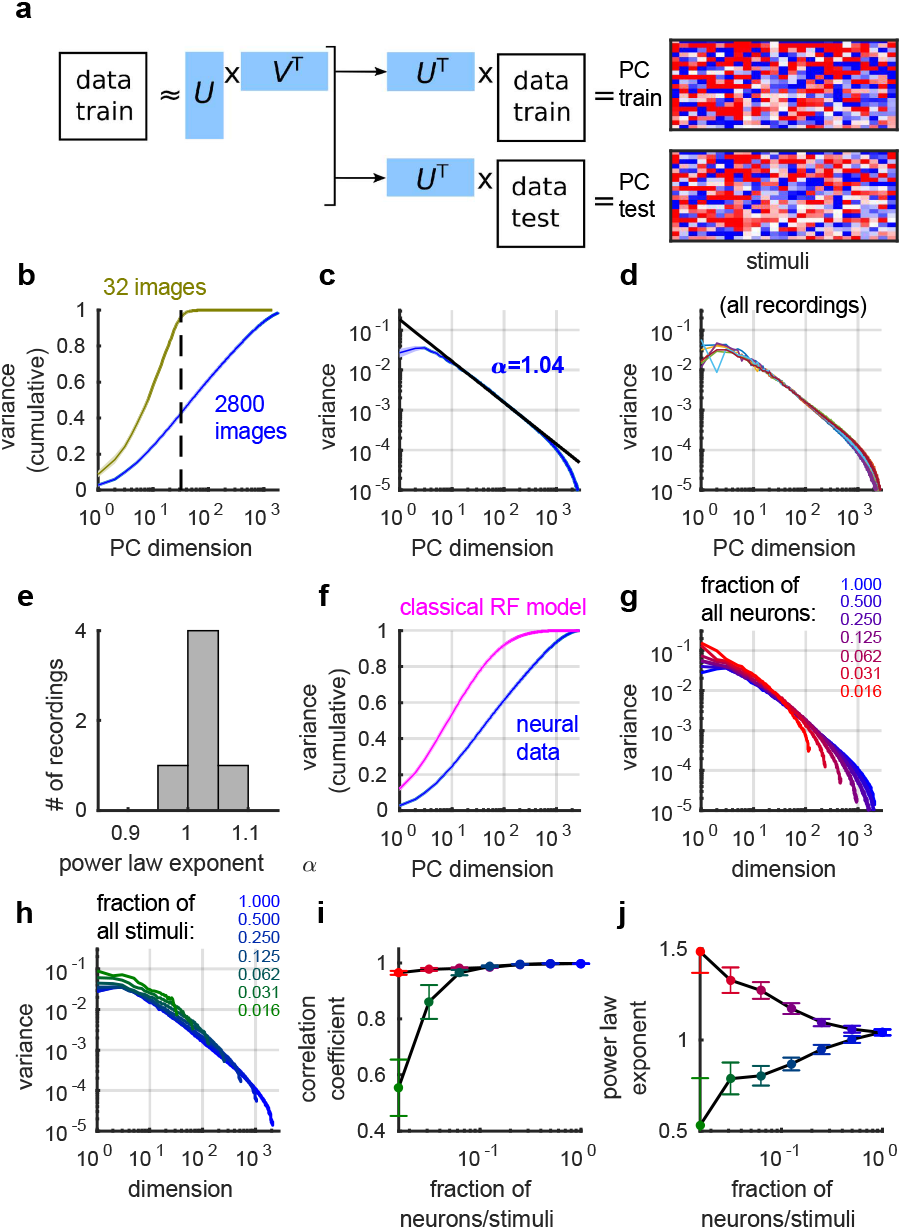
Visual cortical responses are high-dimensional with power-law eigenspectrum. **a**, The eigenspectrum of visual stimulus responses was estimated by cross-validated principal component analysis (PCA), projecting singular vectors from the first repeat onto responses from the second, **b,** Cumulative fraction of variance in planes of increasing dimension, for an ensemble of 2800 stimuli (blue) and for 96 repeats of 32 stimuli. Dashed line indicates 32 dimensions, **c,** Eigenspectrum plotted in descending order of training set singular value for each dimension, averaged across 7 recordings, **d** Eigenspectrum for each recording individually (superimposed), **e,** Histogram of eigenspectrum power law exponent across all recordings, **f,** Cumulative eigenspectrum for a simple/complex Gabor model fit to the data (pink) superimposed on true data (blue), **g,** Eigenspectra computed from random subsets of recorded neurons, fraction indicated by colors, **h,** Same analysis for random subsets of stimuli, **i,** Correlation coefficients for the log-log eigenspectra plotted in **g,h** as a function of fraction analyzed (1 indicates a power law), **j,** Power law exponents of the spectra plotted in **g,h.**

Cross-validated PCA revealed that the neural code in VI is not low-dimensional: visual population responses did not lie on any low-dimensional plane within the space of possible firing patterns. The amount of variance explained continued to increase as further dimensions were included, without saturating at any dimensionality below the maximum possible (Fig. 2b). As a control analysis, we used the same methods to analyze responses to multiple repeats of a set of only 32 images – whose responses must by definition lie on a 31-dimensional plane – and observed a saturation after 31 dimensions.

This analysis revealed an unexpected finding: the fraction of neural variance in planes of successively larger dimensions followed a power law. For natural image responses, the eigenspectrum, i.e. the function summarizing the variance of the *n^th^* principal component, had a magnitude approximately proportional to 1/*n* (1/*n^α^* where *α* = 1.04) (Fig. 2c, see also Extended Data Fig. 7). This power-law did not result from averaging over experiments: analysis of data from each mouse individually revealed power-law behavior in every case (Fig. 2d). While some variability between mice was observed in the scaling exponent of the power law, this exponent had a peak close to 1 across the population (1.04=L·0.02 SE, n = 7 recordings, Fig. 2e). This eigenspectrum reflected correlations between neurons, and was not the consequence of a log-normal distribution of firing rates or signal variance in the population (Extended Data Fig. 8). In addition, this result could not be explained by classical models of visual cortical receptive fields: a model of simple/complex Gabor receptive fields with parameters fit to single cell responses (Fig. 1h) reached 95% variance explained at 147 dimensions (compared to 998 dimensions in the neural data) (Fig. 2f).

The eigenspectrum power-law we observed was not a function of the finite number of neurons and stimuli we recorded and presented, but grew more accurate the more neurons and stimuli were considered (Fig. 2g-j). By repeating the analyses with randomly-chosen subsets of neurons or stimuli, we found that the correlation coefficient of log-variance with log-dimension grew closer to 1 with increasing numbers of neurons or stimuli (Fig. 2i). Furthermore, the exponent of the power law converged towards 1 with increasing numbers of neurons or stimuli (Fig. 2j). We thus infer that the power law observed in our recordings does not simply reflect how the particular set of neurons we recorded responds to the particular set of stimuli we happened to show; instead this power law represents a universal feature of the neural code in mouse VI, that will also govern the response of even larger neuronal populations to a potentially unlimited ensemble of stimuli with similar visual properties.

### Power-law variances do not arise from natural image statistics

Natural images have a power-law structure^31,32^ (Fig. 3a), but this did not cause the neural code’s power-law. To investigate whether the eigenspectrum of the image set could underlie the eigenspectrum seen in neural responses, we removed the image power law by spatially whitening the images, and presented the ensemble of whitened stimuli to 3 out of the 6 mice. Although the power law in the image pixels was now abolished, the power law in the neural responses remained (Fig. 3b). Furthermore, the eigenspectrum of neural responses could not be explained by simple receptive field properties: a simple/complex Gabor model applied to the input images produced eigenspectra that decayed more quickly than the actual responses, and were worse fit by a power-law (p<10^-3^, Wilcoxon rank-sum test on Pearson correlations) for both the original and spatially-whitened images (Fig. 3a,b).

**Fig. 3.**
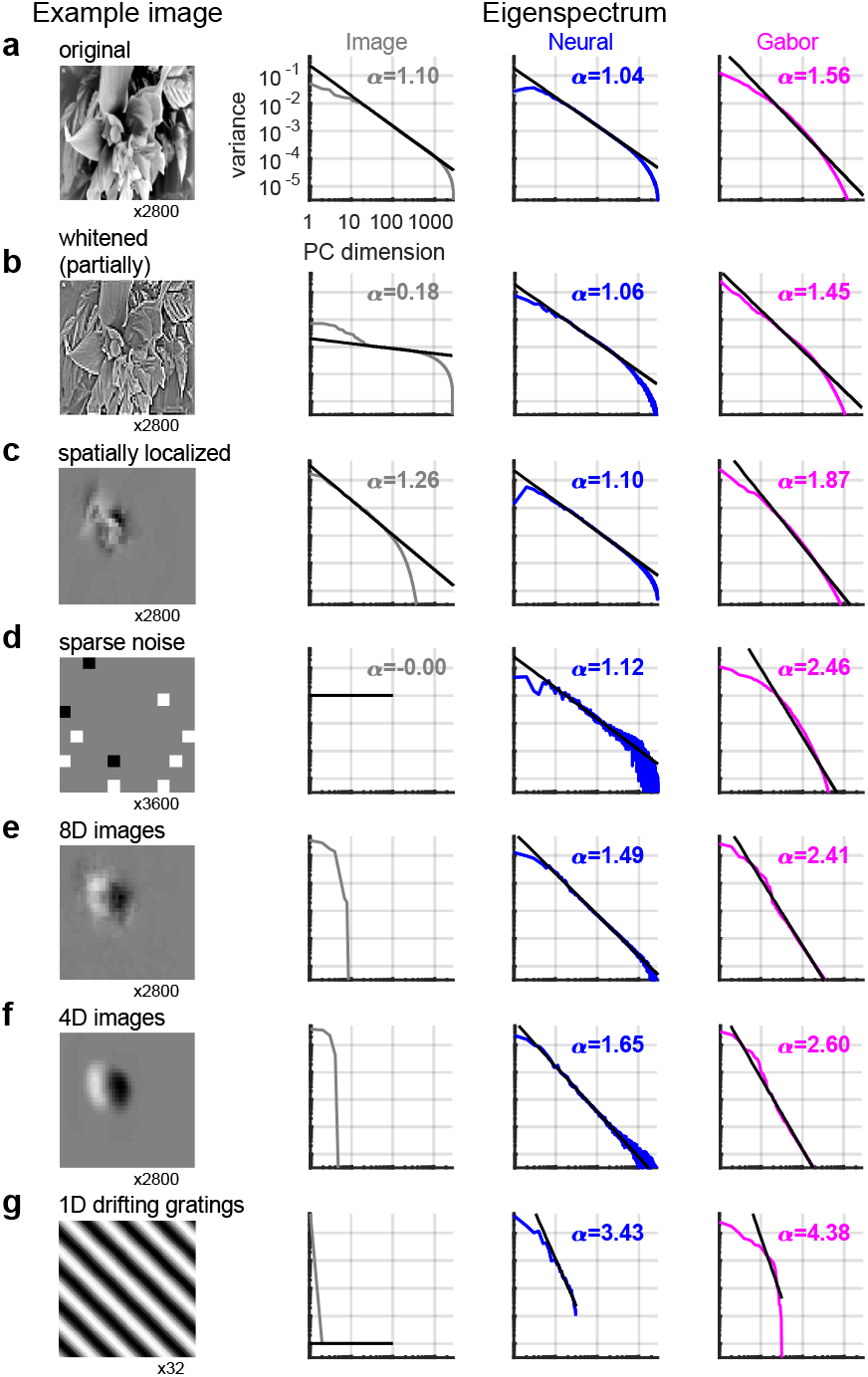
Power law exponent depends on input dimensionality, but not image statistics. **a**, Eigenspectra of natural image pixel intensities (gray), of visual cortical responses to these stimuli (blue), and of a simple/complex cell model’s response to these stimuli (pink), **b,** Same analysis for responses to spatially whitened stimuli, that lack 1/*n* image spectrum, **c,** Same analysis for images windowed over the RF of the recorded population, **d,** Same analysis for sparse noise stimuli, **e,** Same analysis for images projected into 8 dimensions produces a faster eigenspectrum decay with exponent *α*=1.49. **f**, After projecting images to 4 dimensions, *α*=1.65. **g,** Responses to drifting gratings, a one dimensional stimulus ensemble, show yet faster decay with *α*=3.43.

The power law eigenspectra also did not arise from the long-range correlations characteristic of natural images, or indeed from any property unique to natural image stimuli. To investigate the role of long range interactions, we constructed spatially localized image stimuli, in which the region outside the classical RF was replaced by gray. Again, the power law persisted with exponent close to 1 (Fig. 3c). Finally, to investigate whether natural stimuli were in any way required for power-law variances, we showed sparse noise stimuli to the mice (Fig. 3d). Again, we observed a power-law spectrum, with exponent close to 1, although slightly larger than for the natural image stimuli (1.13±0.044 SE, n=3 recordings; *p* = 0.067, Wilcoxon two-sided rank sum test). We conclude that power-law spectra do not reflect neural processing of a property specific to natural images, but also arise in response to multiple types of stimulus ensemble.

### Power-law exponent reflects the dimensionality of the set of inputs

Power law eigenspectra are observed in many scientific domains, and are related to the smoothness of the underlying functions. For example, the Fourier spectrum of a differentiable function of one variable must decay at least as fast as a power law of exponent 1 (see e.g. Ref.^33^). We therefore hypothesized that the variance power law might be related to the smoothness of the set of neural responses. The set of neural population responses to stimuli drawn from a d-dimensional stimulus space lies on a subset - more specifically a manifold - of dimension at most *d*, meaning that the firing rates of all neurons can be described by a nonlinear function of no more than *d* numbers. A manifold is said to be differentiable if these coordinate functions are not just continuous but differentiable. Differentiable manifolds are smoother than non-differentiable ones. Not all manifolds are differentiable; for example, many objects in nature (such as coastlines) are fractal, showing increasing amounts of roughness at finer spatial scales^34^. We showed mathematically that if the set of neural responses is a *d*-dimensional differentiable manifold, then its principal component variances must decay asymptotically faster than any power law with exponent above *α* = 1 + 2/*d* (see Supplementary Information). Conversely, if its eigenvalues asymptotically decay slower than a power law of exponent 1 + 2/*d* then the neural responses cannot lie on a differentiable manifold but must lie on a fractal.

To test the hypothesis that power-law eigenspectra reflected differentiability of the neural response manifold, we presented our mice with stimuli drawn from stimulus ensembles with systematically lower dimension. For a high-dimensional stimulus ensemble such as natural images, *d* will be large so 1 + 2/*d* ≈ 1, which is close to the power law exponent we observed for natural images. However for smaller values of *d*, the power-law must have larger exponents if fractality is to be avoided. We obtained stimulus ensembles of dimensionality *d* = 8 and *d* = 4 by filtering the natural image database, projecting onto a set of basis functions that enforced the required dimensionality (Fig. 3e,f). In addition, we obtained a onedimensional ensemble by showing drifting grating stimuli, which are parametrized by a single number (the grating’s orientation). Consistent with the hypothesis, stimulus sets with d = 8, 4, and 1 yielded power-law scaling of eigenvalues, with exponents of 1.49, 1.65, and 3.43, near the lower bounds of 1.25, 1.50, and 3.00 predicted by the 1 + 2/*d* exponent (Fig. 4a). These results suggest that the neural responses lie on a differentiable manifold, but one that is almost as high-dimensional as possible without becoming fractal. The simulated neural responses obtained from the simple/complex cell model also satisfied the bound, but with higher exponents, suggesting a differentiable, but lower dimensional representation.

**Fig. 4.**
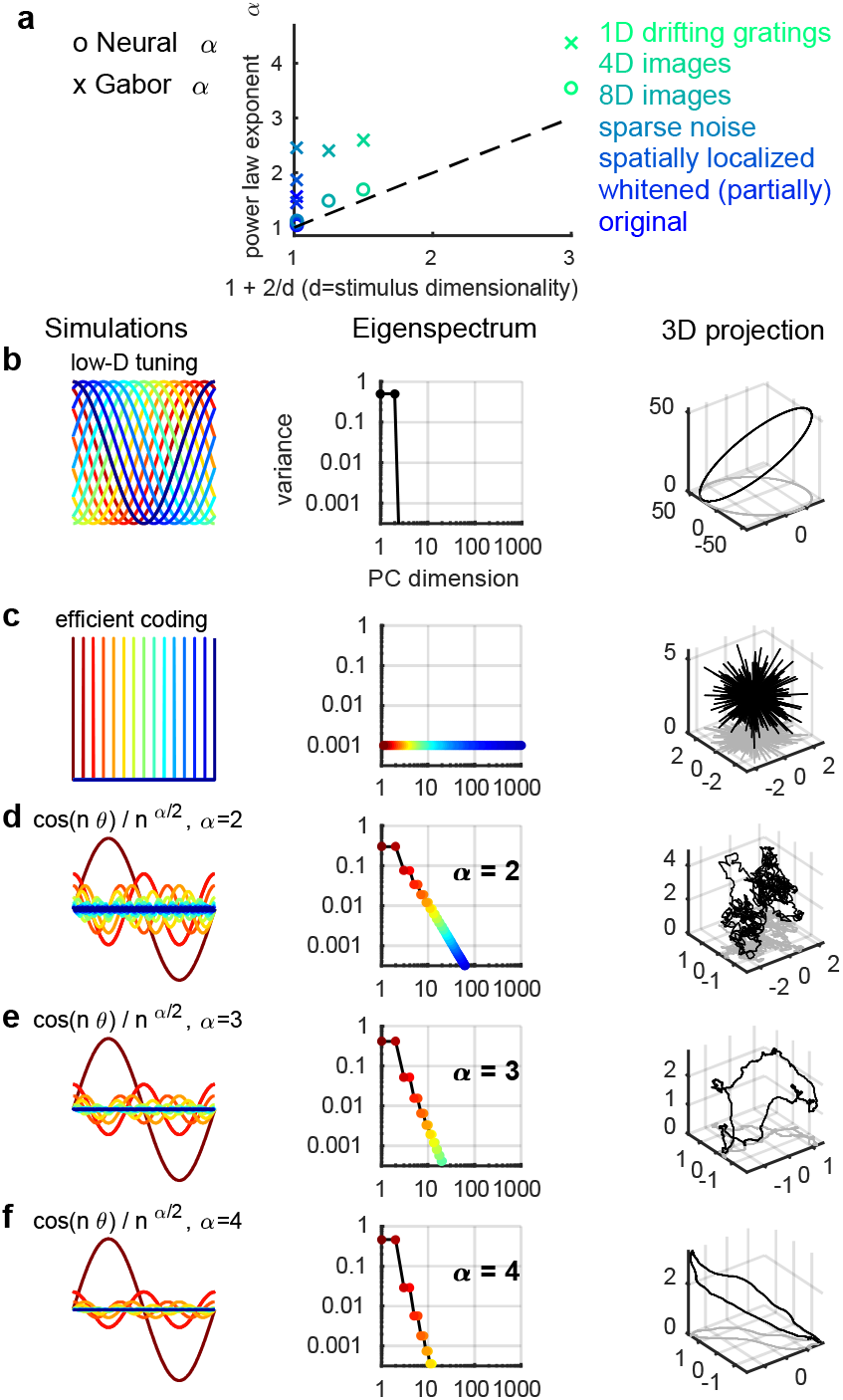
The smoothness of simulated neural activity depends on the eigenspectrum decay. **a**, Summary of power law exponents *α* for neural responses (circles) and Gabor model (crosses), as a function of the dimensionality of the stimulus set *d*. Dashed line: *α* = 1 + 2/*d*, corresponding to the border of fractality. **b-f**, Simulations of neuronal population responses to a 1-dimensional stimulus (x-axis), their eigenspectra, and a random projection of responses in 3D space, **b,** Wide tuning curves, corresponding to a circular neural manifold in a 2-dimensional plane, **c,** Narrow tuning curves corresponding to uncorrelated responses as predicted by the efficient coding hypothesis, **(d-f)** Scale-free tuning curves corresponding to power law variance spectra, with exponents of 2, 3 (the critical value for *d* = 1), or 4.

## Discussion

By analyzing the responses of >10,000 neurons to thousands of image stimuli, we found that visual cortical population responses to stimuli are not constrained to a low dimensional plane. The geometry of the set of stimulus responses was however constrained such that the variance of *n^th^* dimension decayed as a power of *n*, with an exponent of *α* ≈ 1 + 2/*d* where *d* is the dimensionality of the space of sensory inputs. We showed mathematically if the variances decayed slower than this then they could not lie on a differentiable manifold, but must instead lie on a fractal. Nevertheless, these results are not mathematically inevitable: for example, simulating Gabor receptive fields gave responses whose variances decayed faster than this bound. Our experimental results therefore suggest that the eigenspectra of the visual cortical code decay close to the slowest they could, consistent with differentiability of the neural manifold.

To illustrate the geometrical consequences of power-law variance spectra, we simulated neural codes with different eigenspectra (see Supplementary Information), and visualized them through a random projection onto 3D space. The stimulus was 1-dimensional and circular, such as the orientation of a grating, and the population was composed of 1,000 neurons. We first simulated a simple lowdimensional code with two equal variances and all others zero (Fig. 4b). The set of neural responses was by definition constrained to a plane, within which the neural manifold lay on a circle. We then simulated a high-dimensional code in which each neuron responds to a different stimulus, which produces 1,000 equal variances and all others zero (Fig. 4c). These decorrelated responses replicate the population code proposed by the efficient coding hypothesis. Though high-dimensional, when viewed in 3D projection, it appeared as a spiky, discontinuous ball. This manifold is smooth only at scales shorter than the reciprocal of the number of non-zero dimensions (less than 1 degree in this example). For distances greater than this characteristic scale, the code does not respect distances: responses to two stimuli separated by just a few degrees are as different as responses to diametrically opposite stimuli.

Power-law codes show a scale-free geometry, whose de gree of smoothness depends of the exponent *α* (Fig. 4d-f). A power-law code with *α* = 2 (below the critical value of *α* = 3 for a ID stimulus) is a one-dimensional manifold with fractal dimension 2 (see Supplementary Information), which is preserved upon random projection into 3 dimensions (Fig. 4d). The non-differentiable, fractal structure of this manifold is visible by its “fuzzy” appearance, which reveals ever finer details at smaller length scales. A consequence of this fractality is that the fraction of neural variance encoding large-scale stimulus features is outweighed by that encoding ever finer features. At the critical exponent of *α* = 3 (which is equal to 1 + 2/*d* given that *d* = 1), the neural manifold has a fractal dimension of 1 and is on the border of differentiability; this is visible as a geometry that can still represent fine differences between stimuli, but does not let these differences swamp features corresponding to larger stimulus differences (Fig. 4e). A higher exponent leads to a smoother neural manifold, further away from the border of fractality (Fig. 4f).

What are the computational consequences of these different coding geometries, and what advantage might the brain gain from a power-law code with close-to-critical exponent? The efficient coding hypothesis would suggest that information is optimally encoded when responses to different stimuli are as different as possible. However, such codes carry a cost in terms of generalization. This can be seen from an extreme example in which the neural responses to any pair of stimuli are orthogonal: in this case, even stimuli that differ only in tiny details would have completely different responses, and a behavior learned in response to one stimulus could never generalize to another. An example of this behavior can be seen in some neural network architectures that provide a discontinuous set of responses^35^, making them misrepresent “adversarial images” that differ only very slightly from normal images^36^. Differentiability of the neural manifold ensures that similar stimuli will be represented by similar firing patterns, allowing generalization to occur. We suggest that a power-law code of critical exponent therefore represents a balance between efficient, high-dimensional codes, and the ability to generalize at multiple scales.

## Methods

All experimental procedures were conducted according to the UK Animals Scientific Procedures Act (1986). Experiments were performed at University College London under personal and project licenses released by the Home Office following appropriate ethics review.

### Animals and surgery

We used mice bred to express GCaMP6s in excitatory neurons in our recordings: 13 recordings from TetO-GCaMP6s x Emxl-IRES-Cre mice (available as JAX 024742 and JAX 005628); 3 recordings from a Camk2a-tTA, Ai94 GCaMP6s 2tg x Emxl-IRES-Cre mouse (available as JAX 024115 and JAX 005628); and 2 recordings from a Camk2a-tTA, Ai94 GCaMP6s 2tg x Rasgrf-Cre mouse (available as JAX 024115 and JAX 022864). We also used mice bred to express tdTomato in inhibitory neurons (GAD-IRES-Cre x CAG-tdTomato, available as JAX 010802 and JAX 007909) in 17 recordings. In this case, GCaMP6s was expressed virally, and excitatory neurons were identified by lack of tdTomato expression.

Surgical methods were similar to those described elsewhere^19,37^. Briefly, surgeries were performed in adult mice (P35-P125) under isoflurane anesthesia (5% for induction, 0.5-1% during the surgery) in a stereotaxic frame. Before surgery, Rimadyl was administered as a systemic analgesic and lidocaine was administered locally at the surgery site. During the surgery we implanted a head-plate for later head-fixation, and made a craniotomy of 3-4 mm in diameter with a cranial window implant for optical access. In Gad-Cre x tdTomato transgenics, we targeted virus injections (AAV2/l-hSyn-GCaMP6s, University of Pennsylvania Vector Core, 50-200 nl, 1-3 x 1012 GC/ml) to monocular VI (2.1-3.3 mm laterally and 3.5-4.0mm posteriorly from Bregma), using a beveled micropipette and a Nanoject II injector (Drummond Scientific Company, Broomall, PA 1) attached to a stereotaxic micromanipulator. To obtain large fields of view for imaging, we typically performed 4-8 injections at nearby locations, at multiple depths (~500 *μ*m and ~200 μm). Rimadyl was then used as a post-operative analgesic for three days, delivered to the mice via their drinking water.

### Data acquisition

We used a 2-photon microscope (Bergamo II multiphoton imaging microscope, Thorlabs, Germany) to record neural activity, and Scanlmage^38^ for data acquisition, obtaining 10622 ± 1690 (standard deviation) neurons in the recordings. The recordings were performed using multiplane acquisition controlled by a resonance scanner, with planes spaced 30-35 *μ*m apart in depth. Ten or twelve planes were acquired simultaneously at a scan rate of 3 or 2.5 Hz. To verify that this low scan rate did not compromise estimation of neuronal responses, in a subset of experiments we recorded in a single plane imaging configuration (30 Hz frame rate), and downsampled in time by a factor of 12. The downsampled traces contained on average 33.9±5.6% less stimulus-related variance. Simulations (Extended Data Fig. 6) indicated that this change in variance would have no effect on the measured eigenspectra.

The mice were free to run on an air-floating ball and were surrounded by three computer monitors arranged at 90° angles to the left, front and right of the animal, so that the animal’s head was approximately in the geometric center of the setup.

For each mouse, recordings were made over multiple days, always returning to the same field of view (in one mouse, two fields of view were used). For each mouse, a field of view was selected on the first recording day such that 10,000 neurons could be observed, with clear calcium transients and a retinotopic location (identified by neuropil fluorescence) localized on the monitors. If more than one potential field of view satisfied these criteria, we chose either a horizontally and vertically central retinotopic location, or a lateral retinotopic location, at 90° from the center, but still centered vertically. The retinotopic location of the field of view (central or lateral) was unrelated to variance spectra. We also did not observe significant differences between recordings obtained from different modes of GCaMP expression. Thus, we pooled data over all conditions.

### Visual stimuli

Image stimuli were selected from the ImageNet database^20^, from ethologically-relevant categories: “birds”, “cat”, “flowers”, “hamster”, “holes”, “insects”, “mice”, “mushrooms”, “nests”, “pellets”, “snakes”, “wildcat”. Images were chosen manually to ensure that less than 50% of the image was a uniform background, and to contain a mixture of low an high spatial frequencies. Each stimulus consisted of a randomly-chosen image replicated across the three screens after rotating and/or mirroring the image up/down. Stimuli were presented for 0.5 sec, alternating with a gray-screen inter-stimulus interval lasting a random time between 0.3 and 1.1s. For the main experiments, 2800 stimuli were presented twice in the same order each time. Additionally, in a subset of mice (4 out of 6), we presented a smaller set of 32 or 112 images, presented in a randomized order between 32 and 96 times, to enable more accurate estimation of trial-averaged responses.

We also presented partially spatially whitened versions of the 2800 natural images. To compute spatially whitened images, we first computed the two-dimensional Fourier spectrum for each image, and averaged the spectra across images. We then whitened each image in the frequency domain by dividing its Fourier transform by the averaged Fourier spectrum across all images. The rescaled Fourier transform of the image was transformed back into the pixel domain by computing its inverse two-dimensional Fourier transform and retaining the real part. Each image was then intensity-scaled to have similar mean and standard deviation pixel values as the original.

The eight- and four-dimensional stimuli were constructed using a reduced-rank regression model. We first used reduced rank regression to predict the neuronal population responses *R* from the natural images *I* (*N*_pixels_ by *N*_stimuli_) via a *d*-dimensional bottleneck :

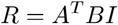

where *A* is a matrix of size *d* by *N*_neurons_ and *B* is a matrix of size *d* by *N*_pixels_. The dimensionality *d* was either eight or four depending on the set of stimuli being constructed. The columns of *B* represent the image dimensions which linearly explain the most variance in the neural population responses. The stimuli were the original 2800 natural images projected onto the reduced-rank subspace *B*: *I*_low-D_ = *B*^⊤^ *BI*.

In addition to natural image stimuli, we also presented drifting gratings and sparse noise. Gratings of 32 directions, spaced evenly at 15° intervals were presented 96 times each, lasting 0.5 sec each, and with a gray-screen inter-stimulus interval between 0.3 and 1.1 s. They were full-field stimuli (all three monitors) and their spatial frequency was 0.05 cycles per degree.

The sparse noise stimuli consisted of uncorrelated squares of size 5° of magnitude ±1 or 0. The probability that a square was non-zero was a uniform distribution of 5%. The squares changed their magnitude every 200 ms. The sparse noise played for 20 minutes, consisting of 6000 unique stimuli in total. Then the same sequence of sparse stimuli was repeated.

In some of the sessions, spontaneous activity was recorded for 30 minutes with all monitors showing a gray or black background. In all sessions, there were occasional blank stimuli (1 out of every 20 stimuli in the 2800 natural image stimuli). The activity during these non-stimulus periods was used to project out spontaneous dimensions from the neuronal population responses (see below).

### Calcium imaging processing

Calcium movie data was processed using the Suite2p toolbox^19^, available at www.github.com/cortex-lab/Suite2P.

Briefly, the Suite2p pipeline consists of registration, cell detection, ROI classification, neuropil correction, and spike deconvolution. Movie frames are registered using 2D translation estimated by regularized phase correlation, subpixel interpolation and kriging. To detect regions of interest (ROIs; corresponding to cells), Suite2p clusters correlated pixels, using a low-dimensional decomposition of the data to accelerate processing. The number of ROIs is determined automatically via a threshold on pixel correlations. Finally, ROIs were classified as somatic or non-somatic using a classifier trained on a set of human-curated ROIs. The classifier reached 95% agreement with training data, thus allowing us to skip manual curation for most recordings. For neuropil correction, we used the approach of Ref.^39^, subtracting from each ROI signal the surrounding neuropil signal scaled by a factor of 0.7; all pixels attributed to an ROI (somatic or not) were excluded from the neuropil trace. After neuropil subtraction, we further subtracted a running baseline of the calcium traces with a sliding window of 60 seconds to remove long timescale additive shifts in the signals. Finally, fluorescence transients were estimated using non-negative spike deconvolution^40^ with a fixed timescale of calcium indicator decay of 2 seconds, a method which we found to out-perform others on ground truth data^41^.

All of the processed deconvolved calcium traces are available on figshare^42^ (https://figshare.com/articles/Recordings_of_ten_thousand_neurons_in_visual_cortex_in_response_to_2_800_natural_images/6845348), together with the image stimuli. The code is available on github (https://github.com/MouseLand/stringer-pachitariu-et-al-2018b).

### Data acquisition and processing (electrophysiology)

Neuropixels electrode arrays^43^ were used to record extracellularly from neurons in six mice. The mice were between 8 and 24 weeks old at the time of recording, and were of either gender. The genotypes of the mice were Slcl7a7-Cre;Ai95, Snap25-GCaMP6s, TetO-GCaMP6s;CaMKΠa-tTA, Ai32;Pvalb-Cre (two mice), or Emxl-Cre;CaMKΠa-tTA;Ai94. In some cases, other electrophysiological recordings had been made from other locations in the days preceding the recordings reported here. In all cases, a brief (<1 hour) surgery to implant a steel headplate and 3D-printed plastic recording chamber (12mm diameter) was first performed. Following recovery, mice were acclimated to head-fixation in the recording setup. During head-fixation, mice were seated on a plastic apparatus with forepaws on a rotating rubber wheel (five mice) or were on a styrofoam treadmill and able to run (one mouse). Three 20×l6cm TFT-LCD screens (LG LP097QX1) were positioned around the mouse at right angles at a distance of 10cm, covering a total of 270×78 degrees visual angle. On the day of recording, mice were again briefly anesthetized with isoflurane while eight small craniotomies were made with a dental drill. After several hours of recovery, mice were head-fixed in the setup. Probes had a silver wire soldered onto the reference pad and shorted to ground; these reference wires were connected to a Ag/AgCl wire positioned on the skull. The craniotomies as well as the wire were covered with saline-based agar. The agar was covered with silicone oil to prevent drying. Probes were each mounted on a rod held by an electronically position-able micromanipulator (uMP-4, Sensapex Inc.) and were then advanced through the agar and through the dura. Once electrodes punctured dura, they were advanced slowly (10*μ*m/sec) to their final depth (4 or 5 mm deep). Electrodes were allowed to settle for approximately 15 minutes before starting recording. Recordings were made in external reference mode with LFP gain=250 and AP gain=500, using SpikeGLX software. Data were preprocessed by re-referencing to the common median across all channels.

We spike sorted the data using a modification of Kilosort^44^ that tracks drifting clusters, which we will refer to as Kilosort2. This modification was necessary to obtain an automated algorithm, and the code will be made publicly available at or before the time of publication. Without the modifications, the original Kilosort and similar algorithms can split clusters according to drift of the electrode. Kilosort2 in comparison tracks neurons across drift levels and for longer periods of time (1 hour in our case). To further mitigate the effect of drift, we used a conservative threshold, excluding from further analysis units for which the maximal firing rate was more than twice their minimal firing rates, when the binned spikes were smoothed with a Gaussian-window filter with a standard deviation of 500 seconds. This excluded 20% of the units on average.

### Removal of ongoing activity dimensions

As shown previously^37^, approximately half the variance of visual cortical population activity is unrelated to visual stimuli, but represents behavior-related fluctuations. This ongoing activity continues uninterrupted during stimulus presentations, and overlaps with stimulus responses only along a single dimension. Because the present study is purely focused on sensory responses, we projected out the dimensions corresponding to ongoing activity prior to further analysis. The top 32 dimensions of ongoing activity were found by performing principal component analysis on mean-subtracted population activity recorded during a 30-minute period of gray screen stimuli before or after the image presentations. To remove these dimensions from stimulus responses, the stimulus-driven activity was first z-scored (using the mean and variance of each neuron computed from spontaneous activity), then the projection onto the 32 top spontaneous dimensions was subtracted (Extended Data Fig. 5).

### Receptive field estimation

We estimated the receptive fields of the neurons, either using a reduced-rank regression model or using a simple/complex Gabor model. In both cases, the model was fit to the mean response of each neuron to half of the 2800 images (*I_train_*) over the two repeats. The performance of the model was tested on the mean response of each neuron to the other half of the 2800 images (*I_test_*).

#### Reduced-rank receptive field estimation

To estimate a linear receptive field for each neuron, we used reduced rank regression^45^, a self-regularizing method which allowed us to fit all neurons’ responses to a single repeat of all 2800 image stimuli. Reduced rank regression predicts high-dimensional outputs from high-dimensional inputs through a low-dimensional hidden “bottleneck” representation. We used it with a 25-dimensional hidden representation to predict each neuron’s activity from the image pixel vectors, taking the resulting regressor matrices as the linear receptive fields. These receptive fields explained 11.4±0.7% (SE, n=7 recordings) of the stimulus-related variance on the test set. These were z-scored prior to display in Fig. 1h and Extended Data Fig. 3.

#### Model-based receptive field estimation

To fit classical simple/complex receptive fields to each cell, we simulated the responses of a convolutional grid of Gabor filters to the natural images, and fit each neuron with the the filter response most correlated to its response.

The Gabor cell filters *G*(***x***) were parametrized by a spatial frequency *f*, orientation *θ*, phase *φ*, size *α* and eccentricity *β*. Defining ***u*** and ***υ*** to be unit vectors pointing parallel and perpendicular to the orientation *θ*:

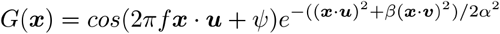

We constructed 12288 Gabor filters, with centers spanning a 9 by 7 grid spaced at 5 pixels, and with parameters *f*, *θ*, *φ*, *α*, and *β* ranging from (0.01,0,0,1,1) to (0.11,157,337,10,2.5)

Simple cell responses were simulated by passing the dot product of the image with the filter through a rectifier function *r*(*x*) = *max*(0,*x*). Complex cell responses were simulated as the root-mean-square response of each unrectified simple cell filter and the same filter with phase *ψ* shifted by 90°. A neuron’s activity was predicted as a linear combination of a simple cell and its complex cell counterpart, with weights estimated by linear regression. Each neuron was assigned to the filter which best predicted its responses to the training images (Extended Data Fig. 4). This simple/complex Gabor model explained 18.4±0.1 % of the stimulus-related variance on the test set.

### Sparseness estimation

The sparseness of neuronal responses was estimated using their responses to 32 natural images. We computed the tuning curve of each neuron by averaging the responses over all 96 repeats. The baseline firing rate of each neuron is computed as the mean firing rate during all periods without visual stimuli (spontaneous activity periods). The standard deviation of the tuning curve is computed for each neuron across stimuli. A cell is responsive to a stimulus if its response to that stimulus is greater than two standard deviations of its tuning curve added to its baseline firing.

### Decoding accuracy from 2,800 stimuli

To decode the stimulus identity from the neural responses, we built a simple nearest neighbor decoder based on correlation. The first stimulus presentation was used as training set while the second presentation was used as test set. We correlated the population responses for a individual stimulus in the test set with the population responses from all stimuli in the training set. The stimulus with the maximum correlation was then assigned as our prediction. We defined the decoding accuracy as the fraction of correctly labelled stimuli.

### Unbiased estimation of signal variance and SNR

At the core of our analysis methods is a method for unbiased estimation of signal variance along any projection of the population activity vector. We first describe this method for the simplest case: estimating the stimulus-related variance of a single neuron. We consider an experiment in which *T* trials are repeated *R* times, with the same stimulus *s_t_* shown on trial *t* in each repeat.

Denote the neuron’s response on repeat *r* of trial *t* as *f_t,r_*. Define *μ_t_* to be neuron’s expected response to stimulus *s_t_*, i.e. the average over a hypothetical infinite number of repetitions (*μ_t_* of course cannot be measured in practice). We can write the neuron’s response as:

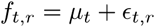

where *ϵ_t,r_* is the trial-to-trial variability, or “noise”. We assume that the noise is independently and identically distributed across repeats of a single stimulus; this condition can be approximately achieved in practice by separating the presentation of the stimulus repeats by tens of minutes to avoid temporally correlated noise. However, we do not assume that the noise has any particular probability distribution, and we allow its distribution and variance to depend on the stimulus.

We would like to estimate the signal variance

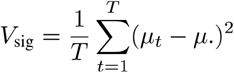

where we employ the usual convention that a dot subscript represents a sample average over the corresponding index. However, the variance computed from a single repeat will also contain an upward bias due to the noise variance. A simple calculation shows that

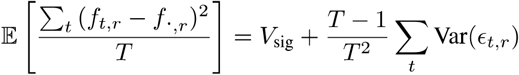

The bias introduced by the noise variance can be reduced, but not eliminated by averaging over repeats. However, we can obtain an unbiased estimate by instead computing the covariance across just two repeats. Indeed, because noise has mean zero and is uncorrelated between repeats,

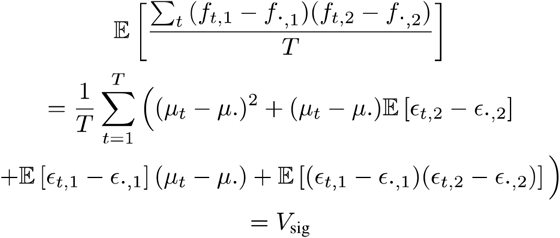

If more than two repeats are available, a the following unbiased estimator can be used:

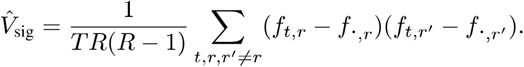

To compute the tuning-related signal-to-noise ratio (SNR; Fig. 1f), we computed the ratio between 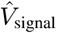 and 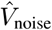 where 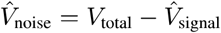 where *V*_total_ is the total variance. This signal-to-noise ratio is non-zero when a neuron has responses to stimuli above its noise baseline.

### Estimating principal component variances

The above method can be extended to obtain an estimate of the principal component variances of the mean population responses to a set of stimuli. We now consider a set of *N* neurons, and define the rate of neuron *n* on repeat *r* of trial *t* as *f_n,t,r_*. The observed firing rate can be written

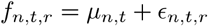

The *N* × *T* matrix ***μ*** is the “signal”, i.e. the average response to each stimulus over a hypothetical infinite number of repeats, and for each repeat *r* the *N* × *T* matrix ***ϵ**_r_* is the “noise”, i.e. the variation from the mean on that repeat. Again, we assume independence between repeats but no other conditions on the distribution of *ϵ_r_*. For example, the noise can be correlated between neurons and have a non-Gaussian distribution that depends on the presented stimulus. Note however that it has zero mean by definition.

We would like to estimate the singular value spectrum of ***μ***. In principle this could be achieved by averaging ***μ*** over multiple repeats, but this would greatly reduce the number of images that can be analyzed. Nevertheless, we can estimate its singular value spectrum from just two repeats by adapting the covariance method of the previous section.

To estimate the singular value spectrum of ***μ*** from only two repeats, we measure the amount of variance in the second repeat that is captured by successive singular vectors of the first. Specifically, we perform a singular value decomposition on the first repeat:

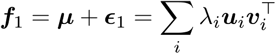

and measure the amount of second-repeat variance explained by the *i^th^* singular vector using the formula

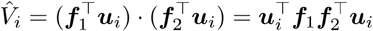

To see why this works, we can expand the firing rates into signal and noise components:

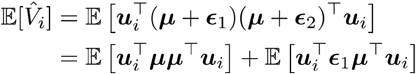

where all terms involving ***ϵ***_2_ have zero expectation due to the statistical independence of ***ϵ***_2_ with ***ϵ***_1_ and ***u***_*i*_, but the second term remains as ***ϵ***_1_ is not independent of ***u**_i_*. In a limit where the singular vectors ***u**_i_* approach the singular vectors of ***μ***, the first term will converge to the singular value spectrum of ***μ***, and the second term will converge to zero (because the variance of ***u**_i_* will also converge to zero). In Supplementary Information section 1, we show mathematically that this convergence will occur, under conditions that hold with good approximation in our recordings.

### Estimation of power-law exponent

We computed the linear fit of the eigenspectrum over the range of 11 to 500 dimensions for all recordings (and model fits) other than the 32-orientation recordings. For the 32-orientation recordings, due to noise and the length of the spectrum, we computed the power-law exponent from 5 to 30. The linear fit was performed in log-log space: the range of log(l 1) to log(500) was regressed onto the log of the eigenspectrum.

## Acknowledgements

We thank Michael Krumin for assistance with the two-photon microscopes, Charu Bai Reddy for surgeries, and Kenneth Falconer and Arthur Gretton for discussions of mathematics.

This research was funded by Wellcome Trust Investigator grants (095668, 095669, 108726, and 205093) and by a grant from the Simons Foundation (SCGB 325512). CS was funded by a four-year Gatsby Foundation PhD studentship. MC holds the GlaxoSmithKline / Fight for Sight Chair in Visual Neuroscience. KDH was funded by the European Research Council (694401). NS was supported by postdoctoral fellowships from the Human Frontier Sciences Program and the Marie Curie Action of the EU (656528). CS and MP are now funded by HHMI Janelia.

## Author Contributions

Conceptualization, C.S., M.P., N.S, M.C. and K.D.H.; Methodology, C.S., M.P., N.S. and K.D.H; Software, C.S. and M.P.; Investigation, C.S., M.P., N.S. and K.D.H; Writing, C.S., M.P., N.S., M.C. and K.D.H; Resources, M.C. and K.D.H. Funding acquisition, M.C. and K.D.H.

---

**Extended Data Fig. 1.**
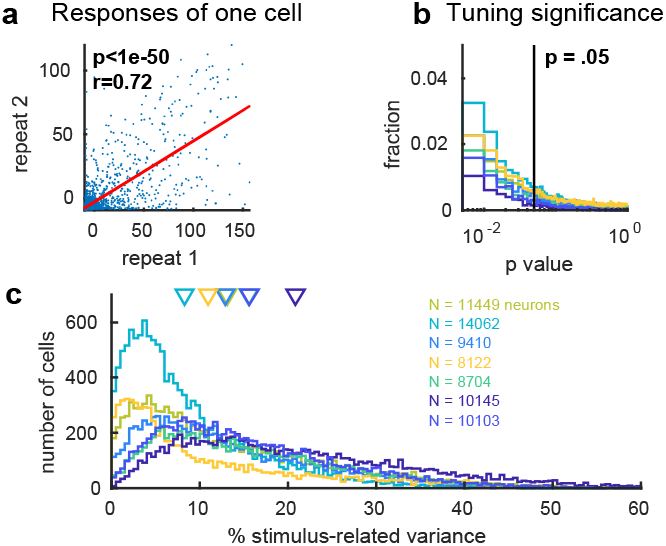
Reliability of single neuron responses. **a**, A single neuron’s response to the first repeat of 2800 stimuli plotted against its responses to the second repeat of the same stimuli, **b,** Histograms of p-values for Pearson correlation of responses on the two repeats. Each line represents a different recording. 81.4 ± 5.1% (SE, n=7 recordings) of cells were significant at p<0.05. **c,** Histogram of the single neuron Pearson correlations across the population.

**Extended Data Fig. 2.**
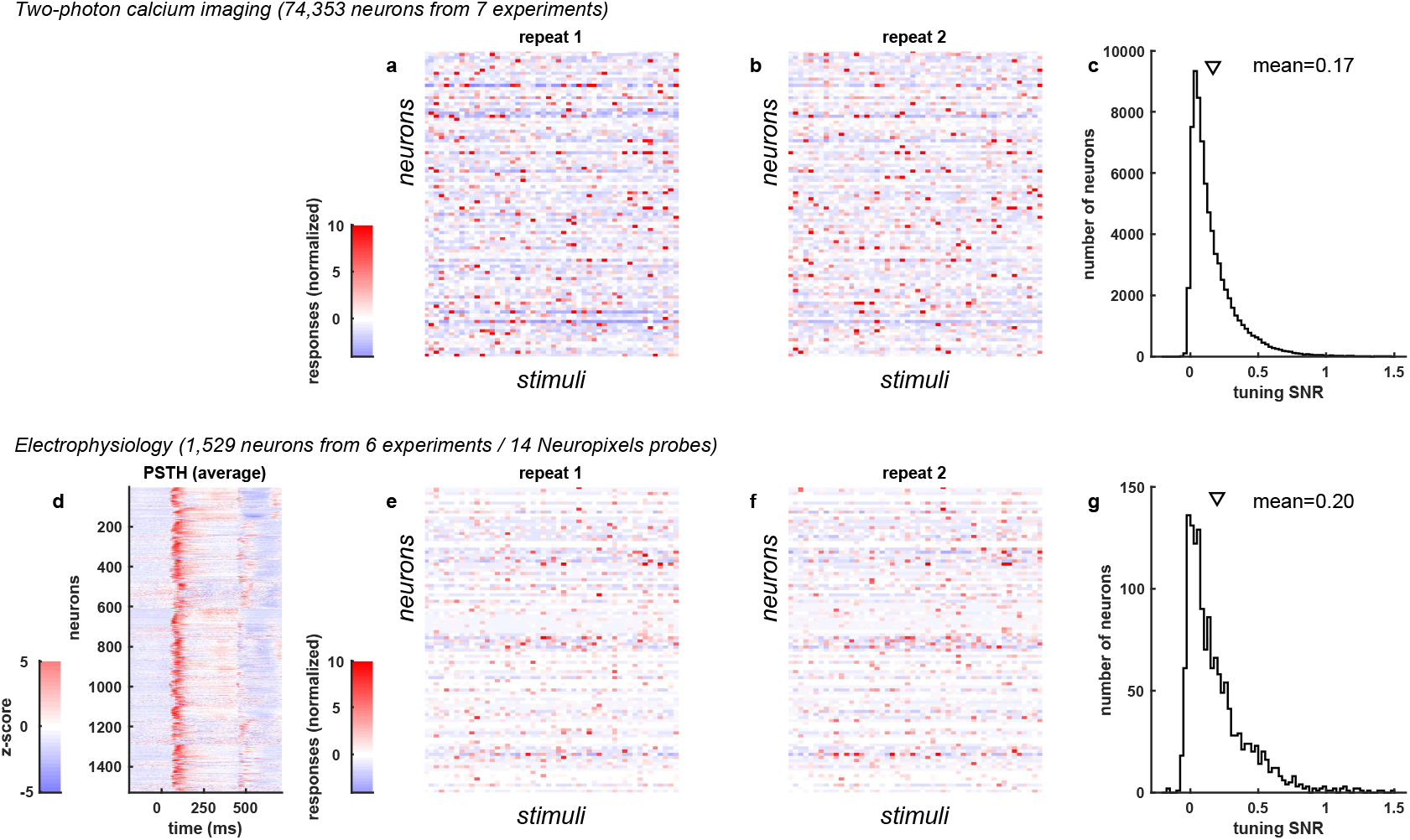
Comparison with electrophysiology. **a**, Single trial responses of 100 neurons recorded by two-photon calcium imaging to 50 stimuli, **b,** Same as (a) for a second presentation of the same stimuli, **c,** Distribution of tuning SNR for 74,353 neurons recorded by two-photon calcium imaging, **d,** Average peri-stimulus time histogram of spikes recorded electrophysiologically in a separate set of experiments. The images shown were a random subset of 700 images out of the total 2,800. **efg** Same as (abc) for the electrophysiologically recorded neurons.

**Extended Data Fig. 3.**
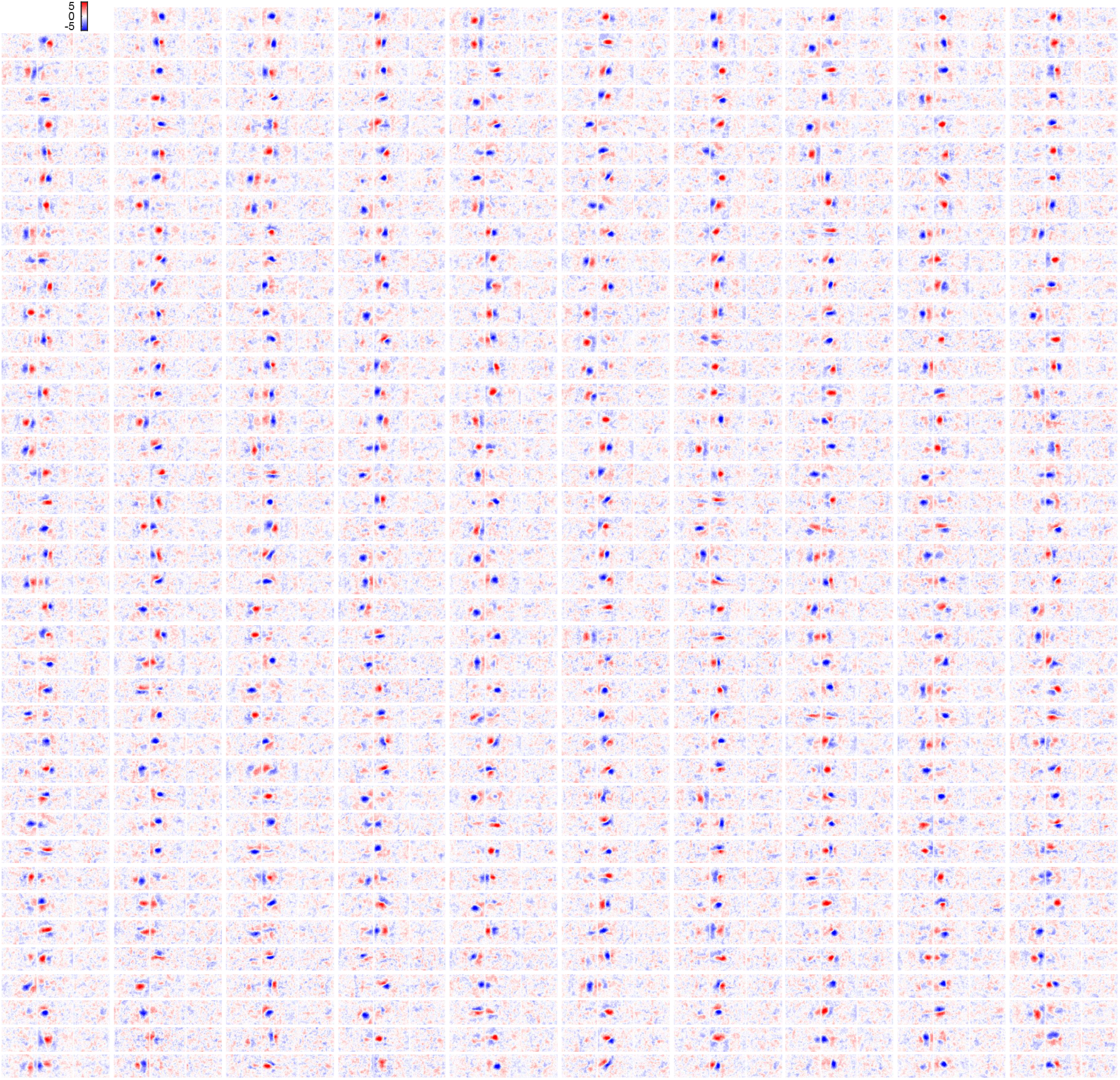
Single neuron receptive fields estimated using reduced-rank regression. 399 randomly chosen neurons’ receptive fields estimated using reduced-rank regression. The receptive field is z-scored for each neuron.

**Extended Data Fig. 4.**
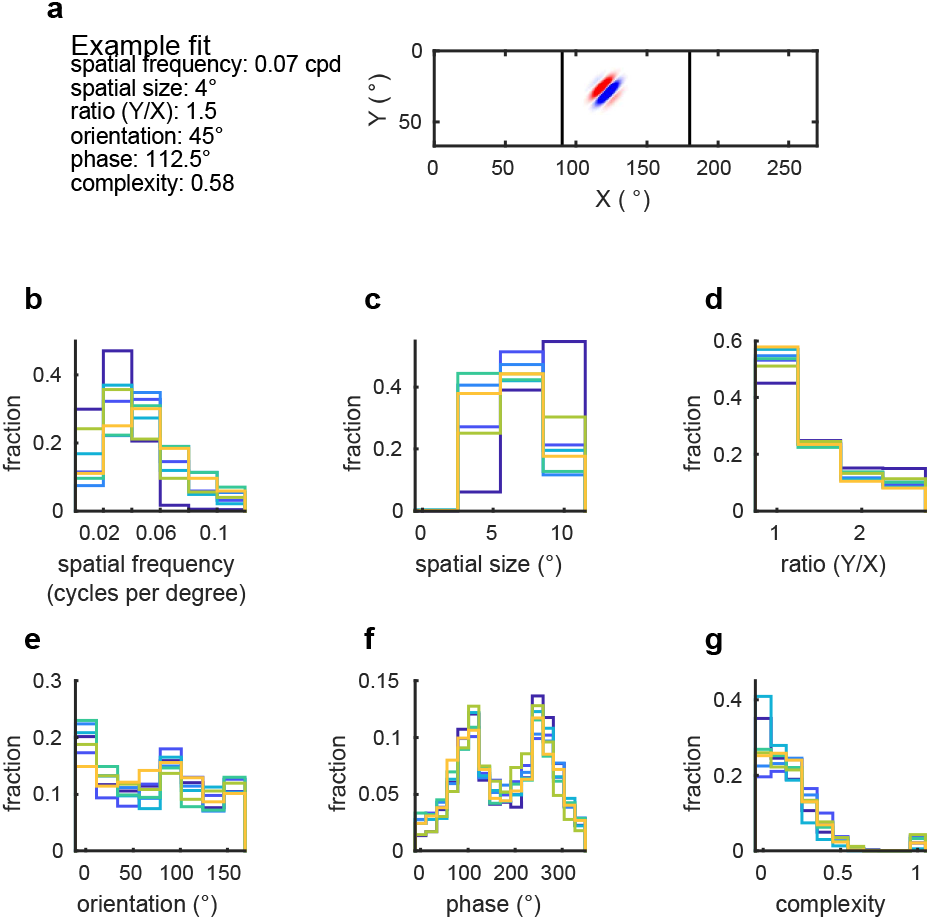
Single neuron receptive field estimation using Gabor models. **a**, An example Gabor fit to a single cell. **b-f** Histograms showing the distribution of model parameters across cells. Each color represents the cells of one recording.

**Extended Data Fig. 5.**
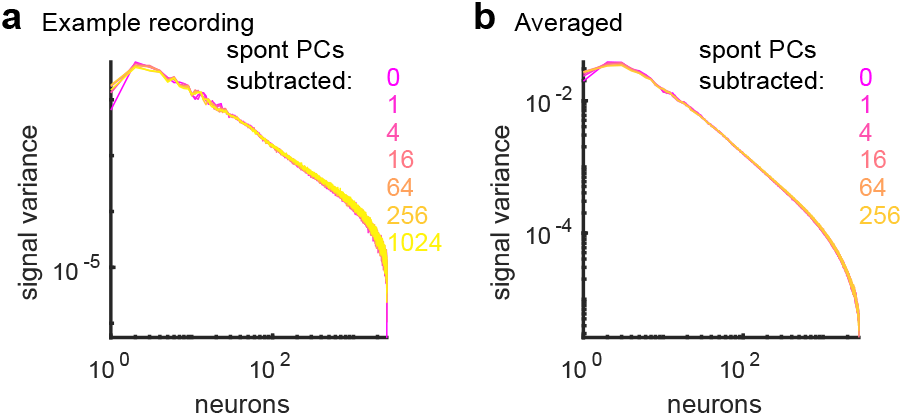
Spontaneous principal component subtraction. **a**, Cross-validated PCA was robust to contamination by stimulus-independent activity. To measure the effects of correlated noise variability on estimated eigenspectra, we examined the effect of projecting out different numbers of noise dimensions (estimated during periods of spontaneous gray-screen) from the responses in an example experiment, **b,** Same analysis, averaged over all recordings. The presence of these noise dimensions made little difference to the estimated signal eigenspectrum other than to slightly reduce estimated eigenvalues in the highest and lowest dimensions. For the main analyses, 32 spontaneous dimensions were subtracted.

**Extended Data Fig. 6.**
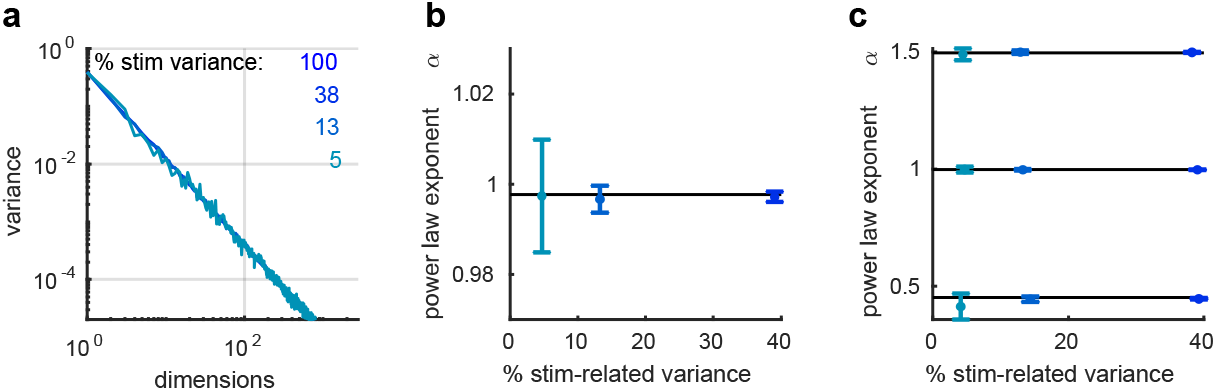
Validating the eigenspectrum estimation method using simulations. **a**, We simulated the responses of 10,000 neurons to 2,800 stimuli with a power spectrum decay of *α* = 1. On top of the responses, we added noise that had the same power-law decay as in the recordings (*α* = 0.70). The simulation was performed for four noise levels, which produced stimulus-related variance ranging from 5% to 100%; for comparison we observed 13.9±1.7% in the neural recordings. In all cases, the *n*^−1^ eigenspectrum was recovered almost exactly (blue curves, color-coded by noise level), **b,** For each level of noise, we simulated 10 different instantiations of additive random noise (with a power-law decay). The average estimated power-law exponent *α* of the simulated responses from each noise instantiation at each noise level is plotted. The error bars are standard deviations across the 10 different noise instantiations, **c** As an additional test, we simulated neural activity where the signal eigenspectrum exponent took different values (*α* = 0.5,1.0,1.5). In all cases, the power law exponent was recovered almost exactly.

**Extended Data Fig. 7.**
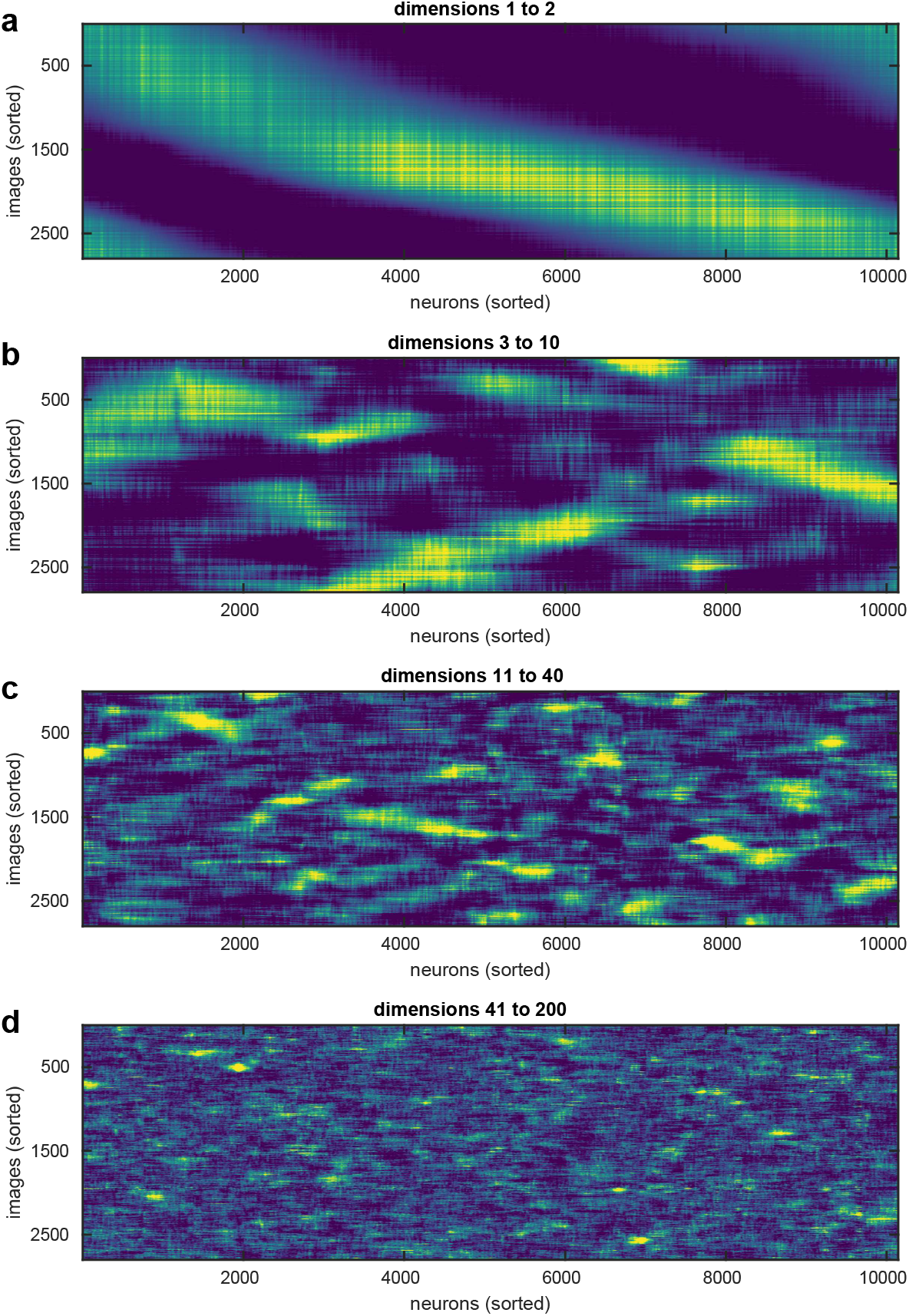
Rasters of neural responses reveal power law scaling. Each plot shows the responses of neurons to the 2,800 natural images projected onto the specified PCs and then sorted along both axes. The sorting placed responses which were more correlated closer to each other. **a,** Dimensions 1-2 reveal a 1D sorting of the neurons and stimuli. **b,** Dimensions 3-10 reveal multidimensional structure which involves subpopulations of neurons. **c,** Dimensions 11-40 reveal smaller dimensions of covariability among neurons. **d,** Dimensions 41-200 reveal even smaller dimensions of covariability.

**Extended Data Fig. 8.**
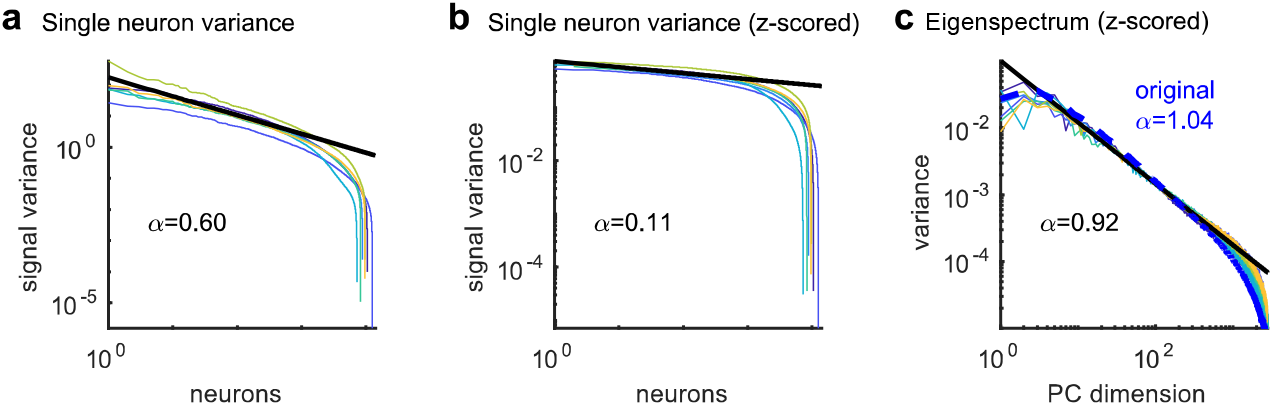
Power law scaling reflects correlation structure, not single-neuron statistics. **a**, The signal variance of each neuron’s responses are sorted in descending order; they approximately follow a power law with a decay exponent of *α* = 0.59. **b,** Same plot after z-scoring the recorded traces to equalize firing rates between cells; the distribution of single-neuron variance has become nearly flat, **c** PC eigenspectra for z-scored data. Each colored line represents a different recording. Dashed blue shows the average eigenspectrum from the original, non-z-scored responses. The fact that the eigenspectrum power-law is unaffected by equalizing firing rates, while the distribution of single cell signal variance is altered, indicates that the power law arises from correlations between cells rather than from the distribution of firing rates or signal variance across cells.

**Extended Data Fig. 9.**
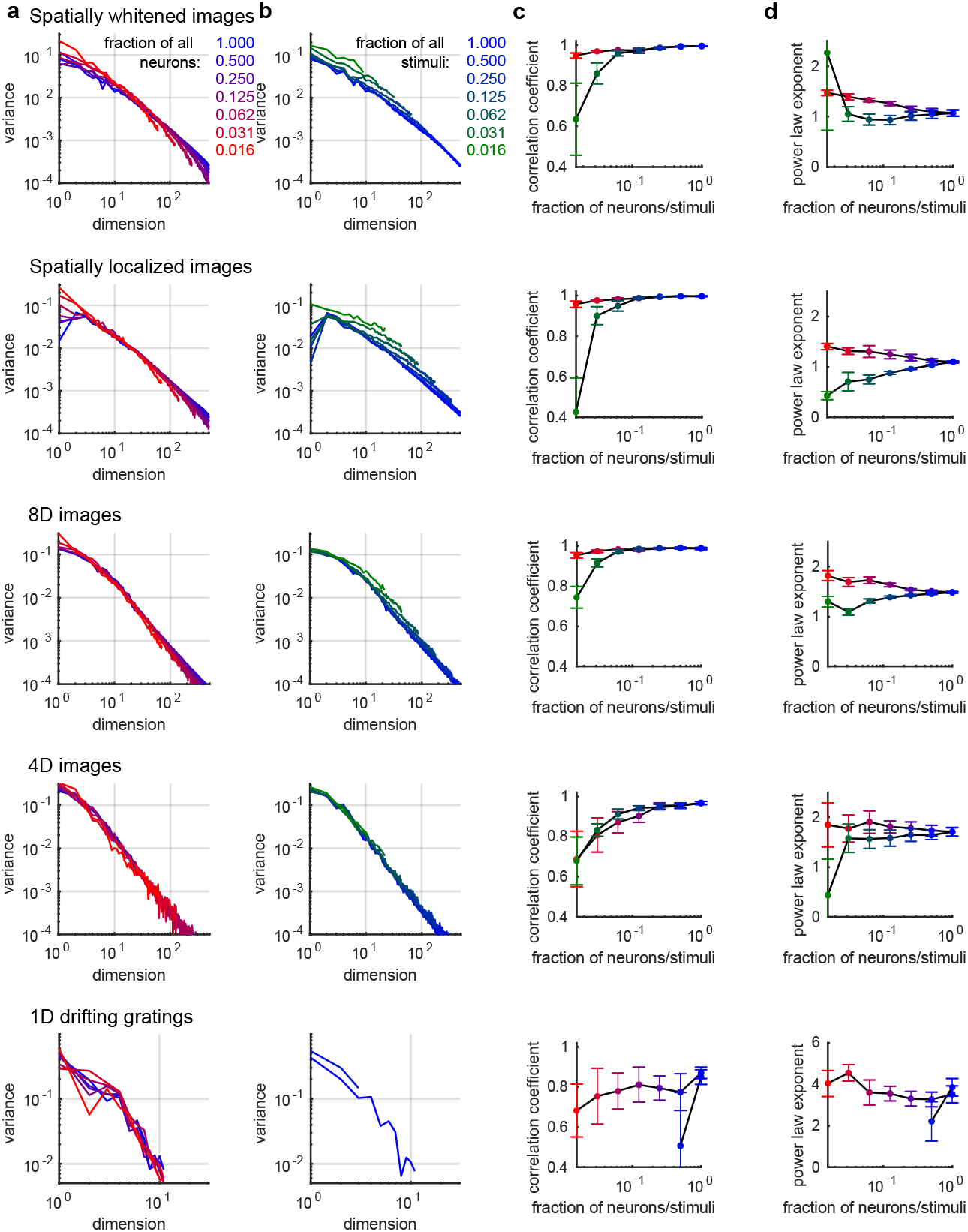
Power law scaling grows more accurate for increasing numbers of neurons and stimuli, for all stimulus ensembles. **a,** Eigenspectra estimated from a random subset of the recorded neurons (color-coded by fraction of neurons retained, **b,** Eigenspectra estimated from a random subset of stimuli, color-coded by fraction of stimuli retained, **c,** Correlation coefficient of the spectra plotted in **a,b. d,** Power law exponent of the spectra plotted in **a,b**. Each row corresponds to a different ensemble of visual stimuli.

## Supplementary Information

The supplementary information contains two sections. In section 1, we show mathematically why the singular vectors estimated from one repeat of all stimuli will be close to those of the noise-free data, in conditions that in our recordings to good approximation. In section 2, we prove that the eigenspectrum of a differentiable manifold of dimension *d* cannot decay slower than a power law of exponent *α* = 1 + 2/*d*.

### 1 The limit of many cells and stimuli

We define three conditions that are sufficient for the estimated singular vectors of the recorded data ***f*** to converge to the singular vectors of the noise-free responses ***μ***. These conditions hold to good approximation in our recordings, as will be discussed at the end of this section.

The first condition is that we record from a sufficiently large number of neurons. We can formalize this requirement using the framework of functional data analysis^1–3^, i.e. the statistical inference of objects that lie in an infinite-dimensional function space. We consider the recorded neurons to have been sampled from a hypothetical infinite population of neurons 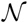, according to a probability measure 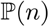, and idealize the firing pattern of visual cortex on trial *t* as a function 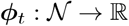, such that *f_n,t_* = *ϕ_t_*(*n*) represents the activity of the neuron *n* on trial *t*. The number of neurons in visual cortex is of course finite, but the use of an idealized infinite population allows mathematical limiting arguments that hold to good approximation. Such limiting approximations occur in all applications of functional data analysis. For example, financial timeseries are composed of a finite number of trades, but functional data analysis provides a way to approximate these very high dimensional discrete data as infinite dimensional continuous functions.

More specifically, we consider the functions *ϕ_t_*(*n*) as lying in the infinite-dimensional Hilbert space 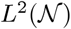, equipped with the inner product 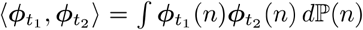. If we now consider a recording of *N* neurons *n*_1_ … *n_N_* randomly sampled from the population 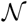, then the law of large numbers implies that as the population size increases, the correlation of their activity on trials *t*_1_ and *t*_2_ converges to the Hilbert space inner product:

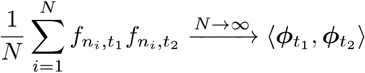

The second condition that we require is to have recorded responses to a sufficient number of stimuli. Again, we formalize this condition by considering the presented stimuli to be drawn at random from a hypothetical infinite ensemble of possible stimuli 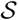, according to a probability measure 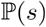. However, the neural activity on trial *t* does not only depend on the stimulus *s_t_* shown on that trial, but also contains a nonsensory (“noise”) component:

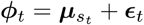

In the functional data analysis framework, correlations are summarized by operators on Hilbert space (the infinite-dimensional generalization of matrices). Thus, we can summarize the correlations of mean responses over the ensemble of all possible stimuli by a “signal correlation operator” 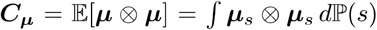. Similarly we can summarize the correlations in the sensory-independent response by a “noise correlation operator” 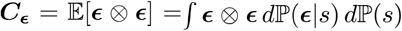. Note we have not assumed that the noise is independent of the stimulus *s*; however, because 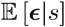 is by definition zero, the noise will be uncorrelated with the signal: 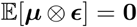, and the total correlation will be the sum of the signal and noise correlations: 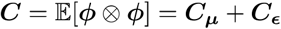.

Now consider a recording of responses on *T* trials, from stimuli *s*_1_ … *s_T_* sampled at random from 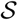, and denote the total correlation measured on these trials as 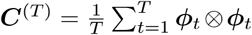. If we write the *i^th^* eigenvector of ***C***^(*T*)^ as 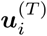, then standard arguments imply that

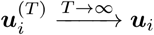

where ***u**_i_* denotes the *i^th^* eigenvector of the total covariance operator ***C***, and convergence should be understood as modulo orthogonal transformation of any eigenvectors with a common eigenvalue (see for example Ref.^2^, theorem 9.1.1).

The third condition we require is that in the limit of a large number of neurons, the noise becomes orthogonal to the stimulus. Although this condition might be unexpected based on low-dimensional intuition, vectors in high dimensional space are orthogonal with high probability; furthermore, an analysis of visual cortical data collected similarly to that analyzed here (Ref.^4^, discussed further below) showed that the spaces spanned by non-visual and visually-evoked activity overlap in only one dimension. We can therefore assume to good approximation that the subspaces of 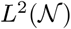 spanned by mean responses ***μ**_s_* and non-visual variability *ϵ_t_* are orthogonal. Denoting the *i^th^* eigenvectors of *C_μ_* and ***C**_ϵ_* as ***u**_ϵ,i_* and ***u**_ϵ,i_* respectively, this orthogonality implies that ***C**_μ_u_ϵ,i_* = ***C_ϵ_u_μ,i_*** = 0. Thus, the set of eigenvectors of the total covariance operator ***C*** = ***C_μ_*** + ***C_ϵ_*** is the union of the sets of eigenvectors ***u**_μ,i_* and ***u**_μ,i_* and the empirical eigenvectors computed from repeat 1 will converge to these for sufficient neurons and stimuli. To determine the results of our cross-validation method in the limit of a large number of neurons and stimuli, it therefore suffices to consider the results for the vectors ***u**_μ,i_* and ***u**_ϵ,i_*.

For any fixed vector ***u***, independence of noise *ϵ*_1_ and *ϵ*_2_ on the two repeats implies

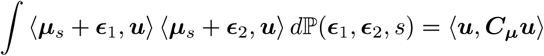

For the signal eigenvectors, therefore

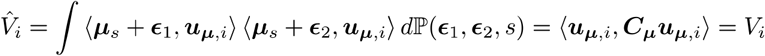

For the noise eigenvectors, we have

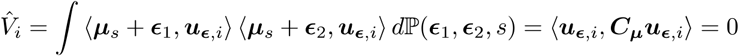

due to the orthogonality of noise and signal in the limit of large neuronal population size. Thus, we conclude that if sufficient neurons and stimuli are recorded, our method should reproduce the signal eigenvalue spectrum, but adding a number of estimated eigenvalues whose expected value is zero, resulting from the noise eigenvectors. Examining the empirical eigenvalue distributions for our neural recordings, we indeed frequently observe a “cliff drop” around the 1000th eigenvalue, after which the eigenvalues fall below the power law (Fig. 2c,d). We suggest this reflects the estimates corresponding to noise eigenvectors. The number of these near-zero estimated eigenvalues is reduced by preprocessing the data to project out the largest non-visual dimensions estimated from spontaneous activity (Extended Data Fig. 5).

Finally, we consider how closely our data obeys the three conditions required for accurate estimation of signal eigenvalues. To test the first two conditions (sufficient neurons and stimuli), we performed a subsetting analysis (Fig. 2g-j). The eigenvalue power law became more accurate the more neurons and stimuli were considered, and the measured exponent appeared to be close to convergence at the numbers of neurons and stimuli used. The third condition, orthogonality of noise and signal spaces, is supported by several observations. Much of what is called neural “noise” is not really random, but encodes nonvisual variables to do with behavioral and cognitive state, a substantial fraction of which can be predicted from videographic analysis of mouse behavior. Analysis of data recorded similarly to that analyzed here^4^ showed that these nonvisual variables are encoded in dimensions almost entirely orthogonal to those encoding sensory stimuli, with the overlap occurring primarily in a single dimension corresponding to population-averaged firing rate. This overlap may have led to underestimation of the first few signal variance components seen in our experimental results, which do indeed dip below the predicted power law (Fig. 2c,d). Not all response variability is predictable from videographic behavior monitoring: for example, “independent noise” that is largely uncorrelated between neurons cannot be predicted videographically. However, in the limit of large numbers of neurons, any sample of this independent noise will become increasingly orthogonal to any predefined vector (including the signal eigenvectors) with probability 1. Finally, another potential violation of the orthogonality condition is multiplicative neural variability: the observation that sensory responses on different trials can represent the same template scaled by different factors^5,6^. However, multiplicative variability will simply scale the expected estimated covariance operator, and will therefore affect neither its eigenvectors nor the estimated signal eigenvalues. We therefore conclude that all three conditions required for eigenvector convergence hold in our data to good approximation.

#### Simulations

To further verify these mathematical arguments, we performed simulations where ***μ*** had a 1/*n^α^* power-law distribution of singular values with varying *α* levels. The simulation was a multiplication of two matrices Λ_*α*_ and *Q*_fix_ where Λ_*α*_ is a matrix of all zeros other than the diagonal:

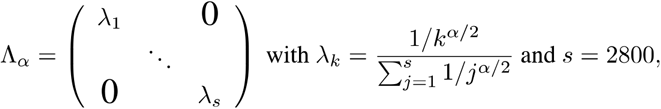

and *Q*_fix_ is

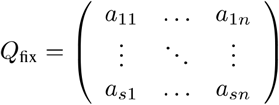

where *a_ij_* is a random number drawn from a Gaussian distribution with mean zero and standard deviation 1, and *n* = 10000 is the number of neurons. We added noise to the simulations which had a 1/*n^β^* power-law distribution where *β* = 0.71 (the noise spectrum in the recordings). The noise was a multiplication of Λ^*β*^ and *Q_t_*, where *Q_t_* is a different instantiation of random Gaussian noise for each trial *t*. The resulting responses from two trials are defined as

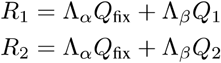

We then computed the eigenspectrum from these two trials of responses using the cross-validated PCA technique described above. We found that the estimated spectrum indeed represented an unbiased estimate of the true spectrum of 1/*n^α^* for varying power-law exponents of the eigenspectrum (Extended Data Fig. 6).

### 2 Mathematical derivation of power-law eigenspectrum bound

In this section we prove that the eigenspectrum of neuronal responses on a differentiable manifold of dimension *d* must decay at least as fast as a power law of exponent *α* = 1 + 2/*d*, which implies that if the variance spectrum decays slower than this the response space must be fractal.

#### Manifolds and Minkowski dimension

There are multiple ways to quantify the dimension of a set, which for some sets do not give the same answer. A manifold is a set that is locally similar to a Euclidean space: in any local region of a *d*-dimensional manifold, each point can be uniquely identified by *d* real-valued coordinates. However, there are two distinct concepts of a manifold, that place different requirements on the coordinate functions. In a *topological manifold*, the coordinate functions are required to be continuous; in a *differentiable manifold*, they are required to be differentiable. Because differentiable functions are always continuous, every differentiable manifold is also a topological manifold. However not all topological manifolds are differentiable, as not all continuous functions are differentiable; indeed, there are functions that are continuous everywhere but differentiable nowhere^7^.

A prime example of topological manifolds that are not differentiable are *fractals*. For example, the coastline of a country can be considered as a one-dimensional topological manifold, but not a differentiable manifold as it shows “rough” structure down to very small scales. Non-differentiability of a manifold can often be revealed by its *fractal dimension*. Fractal dimensions are measures of dimensionality on arbitrary sets, which take the value *d* on a *d*-dimensional differentiable manifold but can exceed *d* if the manifold is not differentiable. The west coast of Britain, for example, has a fractal dimension of approximately 1.25 (Ref.^8^).

There are several ways of formalizing the notion of fractal dimension (See e.g. Refs.^9,10^), all based on the idea that the volume of a *d*-dimensional object with diameter *δ* should scale as *δ^d^*. Here, we will use the *upper Minkowski dimension*. Given a subset *S* of a metric space, we define the the *covering number N_δ_*(*S*) to be the smallest number of spheres of diameter *δ* required to cover *A*. Intuitively, *N_δ_*(*S*) should scale as *δ*^−*d*^ for a space of dimension *d*. The upper Minkowski dimension of *S* is written as 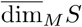, and defined to be inf{*s* : lim sup_*δ*→0_ *N_δ_*(*S*)*δ^s^* = 0}. This means that for any 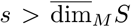 and for any *C* > 0, there exists an *ϵ* > 0 such that for all *δ* < *ϵ*, *N_δ_*(*S*)*δ^s^* < *C*; furthermore 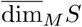 is the smallest number with this property.

This definition involves a limit as a length scale *δ* tends to 0. For a real-world object such as a coastline, it clearly makes no sense to talk about its shape at subatomic scales. Saying a coastline has fractal dimension *d* really means that there is a range of length scales over which *N_δ_*(*S*) scales as *δ*^−*d*^ to good approximation; the limit *δ* → 0 is an idealization that allows theorems to be proved. In the current study, where we consider the dimension of the set of neural responses to visual stimuli, we idealize the visual stimuli as coming from a continuous distribution, even though the monitor has a finite number of possible pixel intensities, and we approximate firing rates as continuous variables even though neurons fire a discrete number of spikes. Furthermore, we idealize firing rate vectors as lying in an infinite-dimensional Hilbert space, which allows limiting arguments about the scaling of covariance eigenvalues.

#### Relating Minkowski dimension to covariance eigenvalues

We now prove our main theorem, which relates the covariance eigenvalues of a random variable supported on a subset of Hilbert space, to that subset’s upper Minkowski dimension.

**Theorem.** *Let X be a random variable supported on a set S of upper Minkowski dimension d inside a separable Hilbert space* 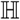, *with* 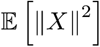. *Write the eigenvalues of Cov*(*X*) *as* λ_1_ ≥ *λ*_2_ ≥ ··· *Then for all s* > *d*, *λ_n_* = *O*(*n*^−1−2/*s*^) *as n* → ∞.

**Proof.** We assume without loss of generality that 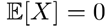. Fix 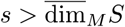 and *C* > 0. It follows from the defintion of the upper Minkowski dimension that there exists *n*_0_ such that for any *n* ≥ *n*_0_, we can cover *S* with *n* balls of diameter at most *Cn*^−1/*s*^. For each point *x* ∈ *S*, let *b*(*x*) be an integer between 1 and *n* identifying a ball in this cover that contains *x*. Define a mean vector for each ball: 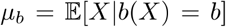. We can decompose the variance of *X* using an ANOVA decomposition:

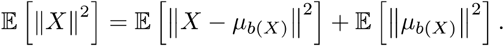

Let *P* be the projection operator onto the *n*-dimensional subspace spanned by mean vectors {*μ_b_* : *b* = 1… *n*}. Then the total variance of *X* in this subspace is at least as big as the variance between means: 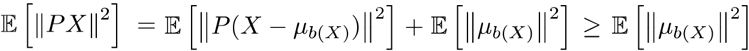. But the variance in any *n*-dimensional subspace cannot exceed the sum of the first *n* covariance eigenvalues: 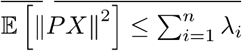 (e.g. Ref.^2^, theorem 7.2.8). Thus,

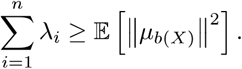

Write the sum of all eigenvalues above *n* as 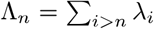. Because the eigenvalues must sum to the total variance, 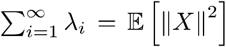, we have 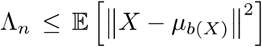. Furthermore, because the balls have diameter at most *Cn*^−1/*s*^, this implies that for all *n* ≥ *n*_0_,

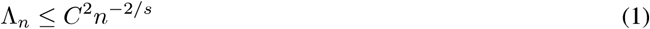

Informally we can see why *λ_n_* should follow an *n*^−1−2/*s*^ power law by differentiating with respect to *n*. Formally, we argue from the convexity of Λ_*n*_. Because the eigenvalues were arranged in non-increasing order, for all *m* ≤ *n* we have 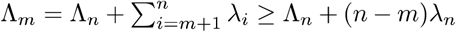. Applying (1) to Λ_*m*_, we have that Λ_*n*_ + (*n* − *m*)*λ_n_* ≤ *C*^2^*m*^−2/*s*^ and since Λ_*n*_ ≥ 0,

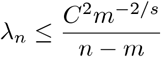

This bound holds for all integer *m* with *n*_0_ ≤ *m* ≤ *n*. To obtain an upper bound for *λ_n_*, we observe that *f_n_*(*m*) = *C*^2^*m*^−2/*s*^/(*n* – *m*) is a convex function of *m* (considered as a real number), with a minimum at *m*_0_ = 2*n*/(*s* + 2), where it takes the value 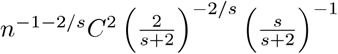. Although *m*_0_ may not be an integer, we can apply the bound to the next largest integer [*m*_0_]. By convexity,

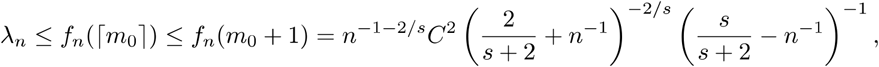

which is *O*(*n*^−1−2/*s*^) for any *s* > 0. Thus, for any number 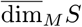,

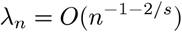

**Corollary.** *If* 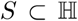 *is a the image of a set of upper Minkowski dimension d under a Lipschitz map, or if S is a d-dimensional compact differentiable submanifold of* 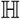, *then for all s* > *d, λ_n_* ~ *O*(*n*^−1−2/*s*^).

**Proof.** In both cases we must show that *S* has upper Minkowski dimension at most *d*. It is a standard result that if *ϕ* is a Lipschitz map, 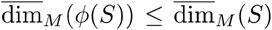, and that if *ϕ* is bi-Lipschitz, 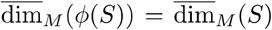 (See e.g. Ref.^10^).

By a *d*-dimensional differentiable submanifold of 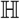 we mean a set 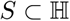, where every point is contained in an open set (in the subspace topology of *S*) that bijects to an open set of 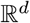 via a chart with a continuous Frechet derivative that is nonzero everywhere. To show such a set has upper Minkowski dimension *d*, observe that each chart must be bi-Lipschitz, and thus each point is contained in an open set of upper Minkowski dimension *d*. By compactness, *S* is therefore covered by finite number of sets of upper Minkowski dimension *d*, and because 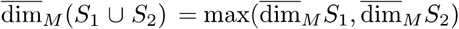 (see e.g. Ref.^10^), 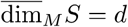.

#### A worked example

We now consider a simple example, in which a 1-dimensional circle is continuously mapped into Hilbert space, with a covariance spectrum of order *n*^−*α*^ by construction. This will show the bounds of the previous section cannot be improved, in the sense that for any *d* ≥ 1, there is a probability distribution supported on an 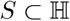 with dim_*M*_ (*S*) = *d* and covariance eigenvalues that decay ~ *n*^−1−2/*d*^. This example is illustrated in Fig. 4d-f.

Parametrize the circle by an angle *θ*, uniformly distributed between 0 and 2*π*. We define a vector *X* (*θ*) in an infinite-dimensional Hilbert space, such that the covariance eigenvalues follow a power law *n*^−*α*^. Specifically, for *n* ≥ 1:

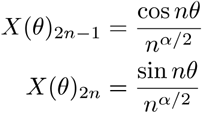

Observe that all dimensions of *X* are uncorrelated; computing their variances shows that *λ*_2*n*−1_ = *λ*_2*n*_ = *n*^−*α*^/2. Also observe that ||*X*(*θ*)||^2^ = *ζ*(*α*), where 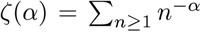 is the Riemann zeta function. Thus ||*X*(*θ*)|| < ∞ if *α* > 1. Furthermore note that 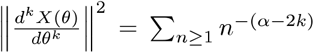. Thus *S* is a topological manifold for *α* > 1, a differentiable manifold for *α* > 3, and *k*-times differentiable manifold for *α* > 1 + 2*k*. However for no value of *α* is *S* an infinitely-differentiable manifold.

We now compute 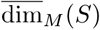. First we compute the distance between two points in *S*. By circular symmetry we may without loss of generality set one of them to 0. We obtain

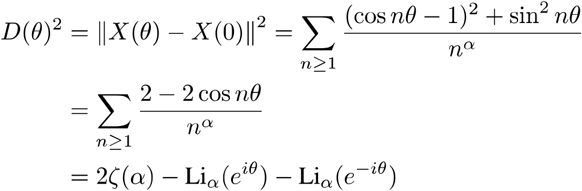

where 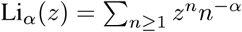 is the polylogarithm function.

To compute 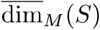 we consider how *D*(*θ*)^2^ depends on *θ* as *θ* → 0. For non-integer *α*, the polylogarithm has a series expansion^11^:

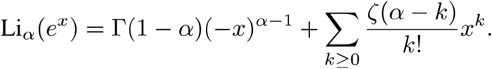

Substituting this in, we obtain

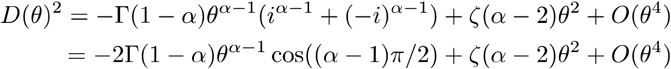

For any value of *θ* we can cover *S* with 2*π*/*θ* balls no smaller than *D*(*θ*). Thus, we have upper Minkowski dimension of *d* if *D*(*θ*) ~ *θ*^1/*d*^ as *θ* → 0. For *α* > 3, the dominant term as *θ* → 0 is *ζ*(*α* – 2)*θ*^2^, so *D*(*θ*) ~ *θ*, and we have 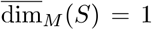. However, for 1 < *α* < 3, the first term of order *θ*^*α*−1^ dominates, so 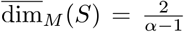, which is greater than 1 indicating a fractal structure. At the critical value of *α* = 3, we can use a Laurent expansion of the the *ζ* and Γ functions to obtain 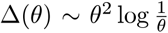. Thus for *α* = 3, the dimension is still 1, even though *X*(*θ*) is not differentiable.

Recall that our theorem predicts that for dimension *d*, we must have *α* ≥ 1 + 2/*d*. This is consistent with the above calculations: *d* = 1 when *α* ≥ 3, and *d* = 2/(*α* – 1) when 1 < *α* < 3. Thus the bound is saturated for 1 < *α* ≤ 3, but loose for *α* > 3.

